# Binding of a pathogen effector to rice Exo70 proteins tethered to the NOI/RIN4 integrated domain of the NLR receptor Pii2 confers immunity against fungi

**DOI:** 10.1101/239400

**Authors:** Koki Fujisaki, Yoshiko Abe, Yu Sugihara, Keiichiro Nemoto, Kazue Ito, Eiko Kanzaki, Kazuya Ishikawa, Mari Iwai, Hiroe Utsushi, Hiromasa Saitoh, Hiroki Takagi, Takumi Takeda, Akira Abe, Shuan Zheng, Aleksandra Bialas, Mark J Banfield, Sophien Kamoun, Ryohei Terauchi

**Author notes:** Co-corresponding authors: Koki Fujisaki and Ryohei Terauchi.

## Abstract

As much as 10% of plant immune receptors from the nucleotide-binding domain leucine-rich repeat (NLR) family carry integrated domains (IDs) that can directly bind pathogen effectors. However, it remains unclear whether direct binding to effectors is a universal feature of ID-containing NLRs given that only a few NLR-IDs have been functionally characterized. Here we show that the rice (*Oryza sativa*) sensor NLR-ID Pii2 confers resistance to strains of the rice blast fungus *Magnaporthe oryzae* that carry the effector AVR-Pii without directly binding this protein. First, we show that AVR-Pii binds the exocyst subunit OsExo70F2 in rice (*Oryza sativa*) to dissociate preformed complexes of OsExo70F2 with host RPM1 INTERACTING PROTEIN4 (RIN4) at the conserved NOI motif, facilitating a possible virulence function. Second, we show that in its resting state, Pii2 binds OsExo70F2 and OsExo70F3, essential components of *Pii*-mediated resistance, through its integrated NOI domain. Remarkably, AVR-Pii binding to OsExo70F2/F3 leads to dissociation of the Pii2–OsExo70F2 and Pii2–OsExo70F3 complexes, destabilization of Pii2, and activation of immunity. These findings support a novel conceptual model in which an NLR-ID monitors alterations of tethered host proteins targeted by pathogen effectors, providing insight into pathogen recognition mechanisms.

**Significance statement:** Plant diseases diminish crop yields by over 20% each year, and deploying resistant crops is the most effective way to combat them. Nucleotide-binding domain leucine-rich repeat (NLR)-type receptors are the major player in plant resistance against pathogens, with a subset of NLRs containing unconventional domains called integrated domains (ID) derived from host proteins. Previous studies suggest that pathogen avirulence (AVR) effectors directly bind or modify NLR-IDs before they are recognized by the host. Here, we reveal that the rice NLR-ID receptor Pii2 indirectly recognizes AVR-Pii when the effector dissociates Pii2 from the host Exo70 proteins tethered to Pii2. We propose a new model of how NLRs can recognize pathogens, expanding our understanding of plant immunity.

## Introduction

Plants have evolved immune receptors to detect and defend themselves against pathogen infections. These receptors include cytosolic nucleotide-binding domain leucine-rich repeat (NLR) proteins that recognize pathogen avirulence (AVR) effectors and activate a complex immune response including a hypersensitive response (HR) leading to localized death of infected cells. Plant NLRs have a modular architecture mostly comprising either an N-terminal coiled-coil (CC) or a Toll/interleukin-1 receptor (TIR) homology domain, followed by nucleotide-binding (NB) and leucine-rich repeat (LRR) domains. Several models have been proposed to explain how NLRs function as singletons, in pairs, or in networks [Adachi et al., 2019]. HOPZ-ACTIVATED RESISTANCE1 (ZAR1), a singleton NLR with N-terminal CC domain of *Arabidopsis* (*Arabidopsis thaliana*), forms a homo-NLR pentamer “resistosome” complex after perception of pathogen AVRs to activate immune responses [Wang et al., 2019]. Several reports have shown that NLR pairs encoded by linked genes (*RESISTANT TO P. SYRINGAE4* [*RPS4*] and RESISTANT TO RALSTONIA SOLANACEARUM1 (*RRS1*) in *Arabidopsis*, *R-GENE ANALOG4* [*RGA4*] and *RGA5* or *Pikp-1* and *Pikp-2* in rice [*Oryza sativa*]) are both required for the recognition of their cognate AVRs [Narusaka et al., 2009; Okuyama et al., 2011; Yuan et al., 2011; Cesari et al., 2014; Zdrzałek et al., 2020]. In these pairs, one NLR functions as a sensor NLR that perceives pathogen effectors, while the other NLR is a so-called helper NLR that activates immune signaling [Baggs et al., 2017; Cesari, 2018].

As much as 10% of all known NLRs contain additional unconventional domains called integrated domains (IDs) that appear to have originated from the host targets of effectors [Cesari et al., 2014; Sarris et al., 2016. The current view is that these NLRs with IDs (NLR-IDs) are baits or decoys that bind pathogen effectors or serve as substrates for effectors to detect invading pathogens. One example is *Arabidopsis* RRS1, an NLR that carries a C-terminal domain with similarity to WRKY transcription factors. RRS1 perceives the bacterial effectors PopP2 from *Ralstonia solanacearum* and AvrRps4 from *Pseudomonas syringae*, which normally target *Arabidopsis* WRKY proteins to enhance virulence. The presence of the WRKY-like domain within RRS1 led to the proposal that this domain targeted by pathogen effectors is integrated into the NLR, enabling recognition of bacterial effectors [Le Roux et al., 2015; Sarris et al., 2015]. Other examples include the rice NLR pairs *Pia* (RGA5 and RGA4) and *Pikp* (Pikp-1 and Pikp-2), which recognize the effectors AVR-Pia/AVR1-CO39 and AVR-PikD of the rice blast fungus, *Magnaporthe oryzae* (syn. *Pyricularia oryzae*), respectively [Okuyama et al., 2011; Kanzaki et al., 2012; Cesari et al. 2013]. RGA5 and Pikp-1 contain a Heavy-Metal-Associated (HMA) domain integrated into different positions [Okuyama et al. 2011; Cesari et al., 2013; Maqbool et al., 2015]. Binding of the integrated HMA domain to the corresponding AVR effectors triggers a HR and immunity against the blast fungus [Cesari et al., 2013; Maqbool et al., 2015; Sugihara et al., 2023]. However, it is unclear whether all NLRs with an ID recognize AVR effectors by direct ID–AVR binding. In this study, we explored the diversity of ID function by analyzing the ID of the rice NLR receptor Pii2, which has a unique feature of pathogen effector recognition.

The *Pii* locus in rice encodes a pair of CC-NLR proteins, Pii1 and Pii2, that detect the *M. oryzae* effector AVR-Pii and mount an effective immune response against the pathogen secreting AVR-Pii [Takagi et al., 2013 and 2017]. Despite repeated attempts using various protein–protein interaction methods, we failed to detect a direct interaction between AVR-Pii and Pii1 or Pii2 [Fujisaki et al., 2015]. Instead, we discovered that AVR-Pii binds to two members of the large EXO70 protein family of rice, OsExo70F2 and OsExo70F3 [Fujisaki et al., 2015; De la Concepcion et al., 2022]; these are putative subunits of the exocyst complex, an evolutionarily conserved vesicle tethering machinery that functions in the last stage of exocytosis [Cvrčková et al., 2012]. Since simultaneous knockdown of *OsExo70F2* and *OsExo70F3* abrogates *Pii*-mediated resistance, we proposed that Pii indirectly recognizes AVR-Pii *via* host Exo70 proteins [Fujisaki et al., 2015].

Here, we show that Pii2 recognition of AVR-Pii follows a mode of NLR perception not previously described for plant pathogens: host Exo70 proteins are tethered to the sensor NLR-ID Pii2, and AVR-Pii binding to Exo70F subunits causes their removal from the NLR-ID, which alters Pii2 status and leads to a HR together with the helper NLR Pii1.

## Results

### Rice Exo70 proteins are required for *Pii*-mediated resistance to the blast fungus

Previously, we showed that simultaneous knockdown of *OsExo70F2* and *OsExo70F3* genes by RNA-interference abrogated rice *Pii*-mediated resistance against *M. oryzae* harboring *AVR-Pii* (Fujisaki et al. 2015). To validate this finding, we used an ethylmethanesulfonate (EMS)-mediated mutant and a CRISPR/Cas9 gene knockout mutant of the *Exo70* genes in the *Pii* rice background. We obtained single mutants of *OsExo70F2* and *OsExo70F3* (*Osexo70f2* and *Osexo70f3*, respectively) and generated a *Osexo70f2 Osexo70f3* double mutant, all in the rice cultivar Hitomebore. Single knockouts of *OsExo70F2* or *OsExo70F3* did not appear to affect *Pii*-mediated resistance, as evidenced by the small lesions seen when these plants were inoculated with the *M. oryzae* strain Sasa2–AVR-Pii (Fig. 1A, 1B; Fig. S1; Vo et al., 2019). By contrast, the *Osexo70f2 Osexo70f3* double knockouts showed no resistance, indicating that OsExo70F2 and OsExo70F3 have redundant functions and each is sufficient for *Pii*-mediated resistance (Fig. 1A, 1B and Fig. S1).

**Fig. 1.**
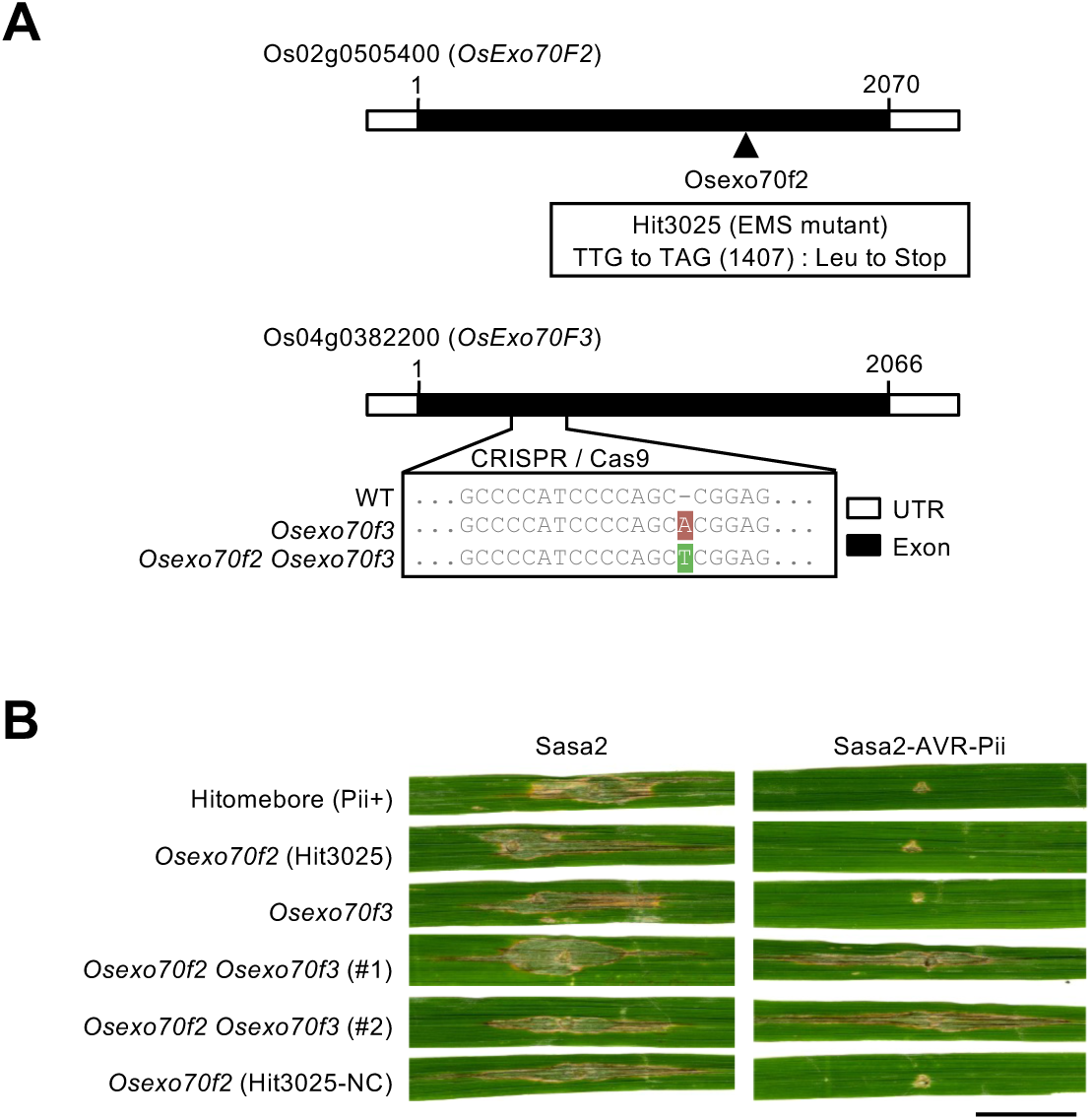
The host *Exo70* genes *OsExo70F2* and *OsExo70F3* are redundantly required in *Pii*-mediated recognition of *AVR-Pii*. (A) Isolation of single and double knockout mutants of *OsExo70F2* and *OsExo70F3*. Hit3025 is a loss-of-function mutant of *OsExo70F2* (*Osexo70f2*) in the rice cultivar Hitomebore (*Pii*+) generated by EMS mutagenesis; *Osexo70f3* is a single knockout mutant of *OsExo70F3* in Hitomebore generated by CRISPR/Cas9-mediated gene editing; *Osexo70f2 Osexo70f3* is a double knockout mutants of *OsExo70F2* and *OsExo70F3* generated by CRISPR/Cas9-mediated gene editing in the Hit3025 background. (B) Effects of *Osexo70f2* and *Osexo70f3* single and double mutations on *Pii*-mediated resistance. Hitomebore and the single and double mutants were inoculated with the rice blast fungus isolate Sasa2 (without *AVR-Pii*) or the transgenic strain Sasa2–AVR-Pii with *AVR-Pii* [Yoshida et al., 2009]. Representative disease lesions on leaves are shown. A transgenic line (Hit3025-NC), which was derived from the same genetic transformation round as *Osexo70f2 Osexo70f3* but with no induced mutation, was used as a negative control. Knockout of *OsExo70F2* and/or *OsExo70F3* is indicated in the left.

### Unlike AVR-Pii, rice Exo70 proteins bind NOI integrated-domain of Pii2

We used protein interaction assays to determine the binding spectrum of Pii2. In a yeast (*Saccharomyces cerevisiae*) two-hybrid (Y2H) assay, we detected an interaction between OsExo70F3 and full-length Pii2 (Figs. S2 and S3A). We delineated the Exo70-interacting region of Pii2 to 103 amino acids in its C terminus (Pii2-CTC for C-terminal half of Pii2-specific CT region; Lee et al., 2009) (Fig. 2A; Fig. S4). In a series of Y2H assays (Fig. 2B, and Fig. S3C), we observed a specific interaction between Pii2-CTC and both OsExo70F2 and OsExo70F3, while AVR-Pii did not interact with any regions of Pii1 or Pii2 (Figs. 2B, S2, S3B, and S5). We confirmed the above interactions using an AlphaScreen (Amplified Luminescent Proximity Homogenous Assay Screen), which relies on the proximity of two potentially interacting proteins to emit fluorescence by acceptor beads (Figs. S6, S7 and S8). Notably, the Pii2-CTC region contains a six-amino-acid stretch showing similarity to the core motif of the nitrate-induced (NOI) domain (PxFGxW) (Fig. 2C and Fig. S9A) [Zhao et al., 2021].

**Fig. 2.**
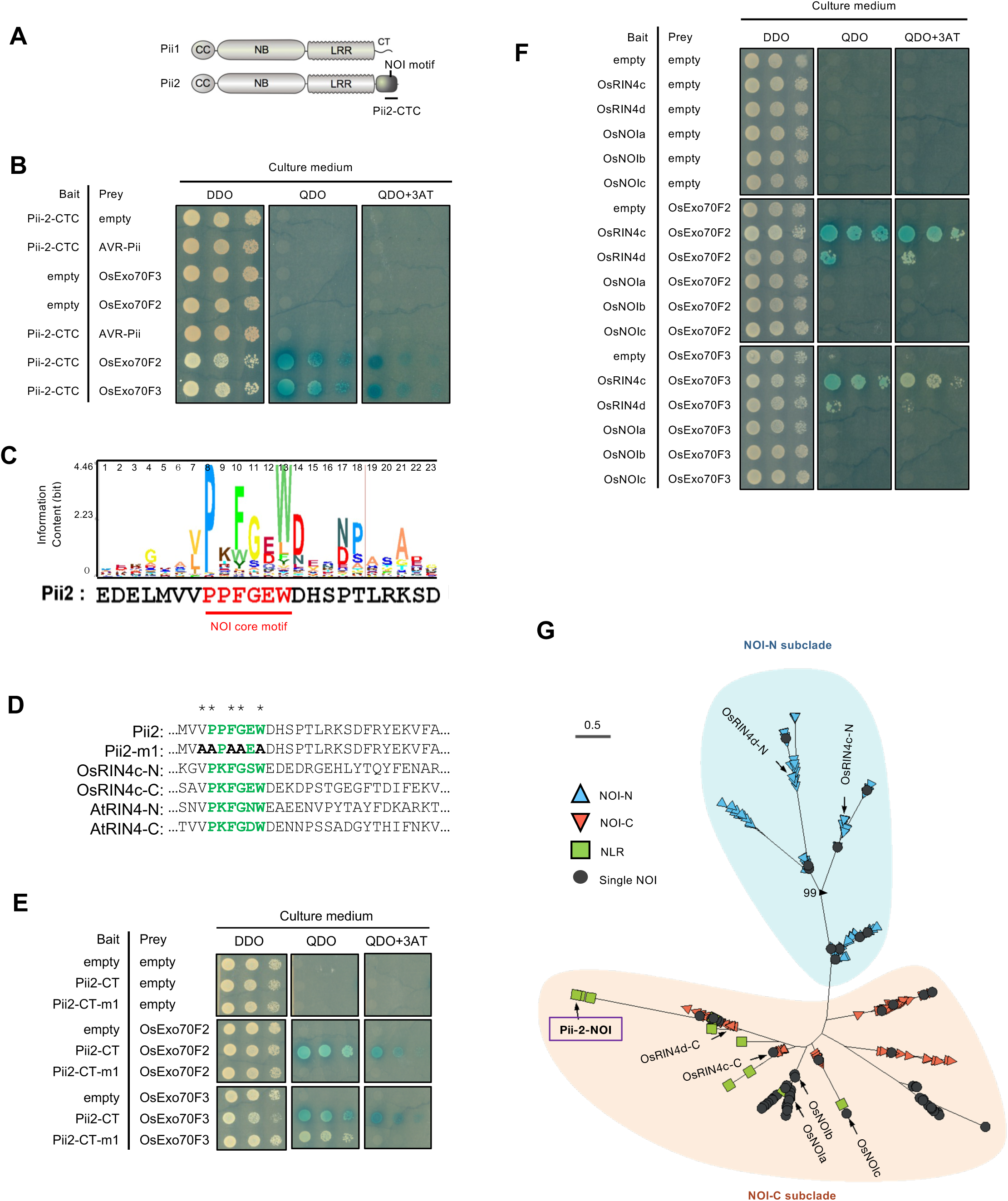
The host Exo70 proteins OsExo70F2 and OsExo70F3, but not the fungal effector AVR-Pii, interact with the NOI–integrated domain (ID) of Pii2, presumably derived from a *RIN4*-like gene. (A) Diagram of Pii1 and Pii2 showing the Pii2 C-terminal fragment (Pii2-CTC) that binds to OsExo70F2 and OsExo70F3. The position of the NOI core motif (PxFGxW) is indicated. (B) Yeast two-hybrid (Y2H) assay testing the interaction of Pii2-CTC with OsExo70F2, OsExo70F3, and AVR-Pii. Ten-fold serial dilution of positive yeast transformants were spotted onto double dropout (DDO) medium as control and on quadruple dropout (QDO) medium and QDO medium + 10 mM 3-amino-1,2,4-triazole (3AT) to test interactions. Empty vector (empty) was used as negative control. For additional data of mapping the interacting regions, see Figs. S2 to S5. (C) Protein logo of the PF05627 domain generated by Skylign (Wheeler et al., 2014). Only the region around the NOI core motif (PxFGxW) is shown. The corresponding amino acid sequence from Pii2 is indicated below. (D) Amino acid sequence alignment of NOI and surrounding sequence in Pii2, the N- and C-terminal regions of rice RIN4c (OsRIN4c-N and OsRIN4c-C) and those of Arabidopsis RIN4 (AtRIN4-N and AtRIN4-C). The conserved amino acid residues are marked by asterisks. The positions of the mutations in the Pii2-m1 mutant are indicated in bold black text. (E) Interaction test between the Pii2-m1 mutant and OsExo70F2 or OsExo70F3 in Y2H. (F) Y2H assay testing the interaction of rice proteins containing a NOI core motif with OsExo70F2 and OsExo70F3. (G) Phylogenetic tree of 737 NOI domains from *Poaceae* species, using only the regions detected as NOI domains using HMMSearch. The phylogenetic tree was reconstructed by the maximum likelihood method using IQ-TREE with 1,000 bootstrap replicates. NLRs were annotated using NLRtracker (Kourelis et al., 2021). When a non-NLR protein contained two NOI domains, the N-terminal and C-terminal NOI domains were labeled as N or C. When a non-NLR protein contained a single NOI domain, the NOI domain was labeled as NOI.

### The RIN4-like NOI motif of Pii2 is required for binding to Exo70

The NOI domain is also present in *Arabidopsis* RPM1-INTERACTING PROTEIN4 (RIN4), which negatively regulates plant defense and is targeted by the bacterial effectors AvrRpt2, AvrB, and AvrRpm1 [Kim et al., 2005]. In fact, the effector AvrRpt2 has protease activity and cleaves RIN4 within its NOI motif [Chisholm et al., 2005]. To investigate the possible importance of the NOI core motif of Pii2, we introduced multiple mutations (Pii2-m1 variant) to replace conserved amino acid residues in this region with alanines (Fig. 2D) and tested the m1 mutant for binding to OsExo70F2 and OsExo70F3. In a Y2H assay and an AlphaScreen, the m1 mutant displayed compromised Pii2 binding to OsExo70F2 and OsExo70F3 (Figs. 2E, S10 and S11). These results highlight the importance of the Pii2 ID NOI core motif in Pii2–OsExo70F2 and Pii2–OsExo70F3 interactions.

### NOI domain of Pii2 has evolved from NOI-C domain of RIN4 proteins

RIN4 proteins have two NOI domains, NOI-N and NOI-C, at their N and C termini, respectively [Afzal et al., 2013]. A BLAST search using *Arabidopsis* RIN4 as query identified six rice RIN4-like proteins (OsRIN4a–f; Fig. S12) and three rice proteins containing one NOI core motif (OsNOIa–c; Fig. S13). *OsRIN4c* and *OsRIN4d* were expressed in rice leaves (Fig. S14), and their protein products interacted with OsExo70F2 and OsExo70F3, unlike the proteins with a single NOI core motif (Fig. 2F and Fig. S15). OsRIN4c showed a strong interaction with OsExo70F2 and OsExo70F3 in Y2H assays (Fig. 2F) and co-immunoprecipitated from protein extracts of *Nicotiana benthamiana* leaves infiltrated with *FLAG-OsRIN4c* and StrepII and HA (SH)-tagged *OsExo70F2* or *OsExo70F3* constructs (Fig. S16). Importantly, mutating the C-terminal NOI (PxFGxW) motif abolished these interactions (Fig. S17). The amino acid sequence of the Pii2 NOI ID indicated that it is more closely related to OsRIN4 NOI-C than to NOI-N (Fig. 2G), suggesting that the NOI core motif of Pii2 is most likely derived from the NOI-C of RIN4-like proteins.

### NOI domain of Pii2 is required for its stability and resistance to blast fungus

To explore the role of Pii2 ID in *Pii*-mediated resistance, we tested the ability of a *Pii2* transgene incorporating the m1 mutation in the sequence encoding the NOI core motif (Fig. 2D) to achieve transgenic rescue. We identified a *pii2* mutant rice line, Hit5882, among 5,600 ethylmethanesulfonate (EMS)-mutated lines of the rice cultivar Hitomebore. The mutant version of Pii2 in Hit5882 carries an amino acid substitution (V371D) in the Pii2 NB-domain (Fig. S9A). When we introduced a transgene comprising the full-length coding sequence of Pii2 cloned in-frame with a FLAG tag sequence (*Pii2FL2*; Fig. S9A) and driven by the cauliflower mosaic virus 35S promoter into Hit5882, Pii2FL2 protein was expressed and accumulated (Fig. 3A) and *Pii*-mediated resistance was restored (Fig. 3B, Fig. S18). By contrast, transgenic Hit5882 plants harboring a 35S:*Pii2FL2-m1* transgene showed low accumulation of Pii2FL2-m1 protein (Fig. 3A) and no *Pii*-mediated resistance (Fig. 3B and Fig. S18). These results suggest that the m1 mutation not only abrogates Pii2 interactions with OsExo70F2 and OsExo70F3 (Fig. 2E and Fig. S10) but also disrupts the stable accumulation of Pii2 *in planta*, demonstrating its serious effects on *Pii*-mediated resistance.

**Fig. 3.**
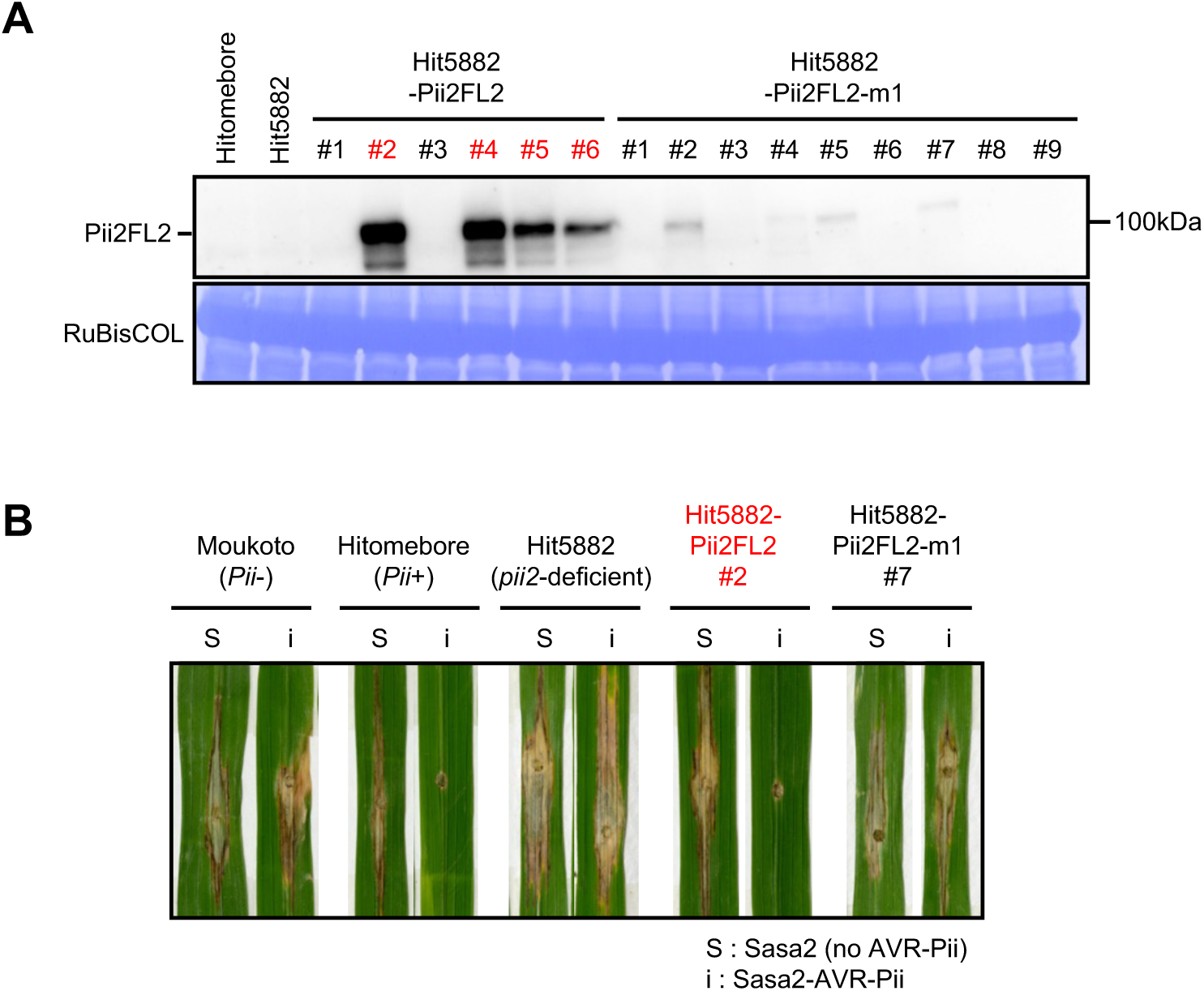
The Pii2 variant carrying the m1 mutation has a low abundance in rice leaves and does not induce resistance against AVR-Pii. (A) Pii2FL2 accumulation in transgenic rice leaves. Pii2FL2 and its m1 variant (Pii2FL2-m1) were detected by immunoblot analysis using anti-FLAG antibodies. CBB-stained RuBisCOL (Rubisco large subunit) are shown below as loading control. (B) Complementation assay by Pii2FL2 or Pii2FL2-m1. Representative data of disease lesions following inoculation of the rice cultivars Moukoto (*Pii*−), Hitomebore (*Pii*+), Hit5882 (a *pii2* mutant in Hitomebore), and Hit5882 transgenic lines expressing Pii2FL2 or Pii2FL2-m1 (Hit5882-Pii2FL2 and Hit5882-Pii2FL2-m1, respectively) with the rice blast fungus strains Sasa2 (S) and Sasa2–AVR-Pii (i). In (A–B), the transgenic lines showing *Pii*-mediated resistance are marked by red letters.

### AVR-Pii co-expression with OsExo70F2/F3 reduces Pii2 protein accumulation

We assessed the link between Pii2 binding to OsExo70F2 and OsExo70F3 and Pii2 accumulation *in vivo*. Accordingly, we co-transfected rice protoplasts with constructs encoding Pii1, Pii2FL2, and OsExo70F2, OsExo70F3, and/or AVR-Pii in various combinations. Although we did not induce a strong HR in this system (Fig. S19), we did notice that co-expression of *Pii2FL2* and *OsExo70F2* or *OsExo70F3* led to enhanced accumulation of Pii2FL2, an effect that was suppressed by co-expression of *AVR-Pii* but not *AVR-Pii^Y64A^* (as negative control) (Fig. 4). By contrast, we did not observe enhanced accumulation of Pii2FL2-m1 when we co-expressed *Pii2FL2-m1* with *OsExo70F2* or *OsExo70F3* (Fig. S20). These results suggest that NOI ID-mediated binding to Exo70 is crucial for Pii function *via* stable accumulation of Pii2 in rice cells.

**Fig. 4.**
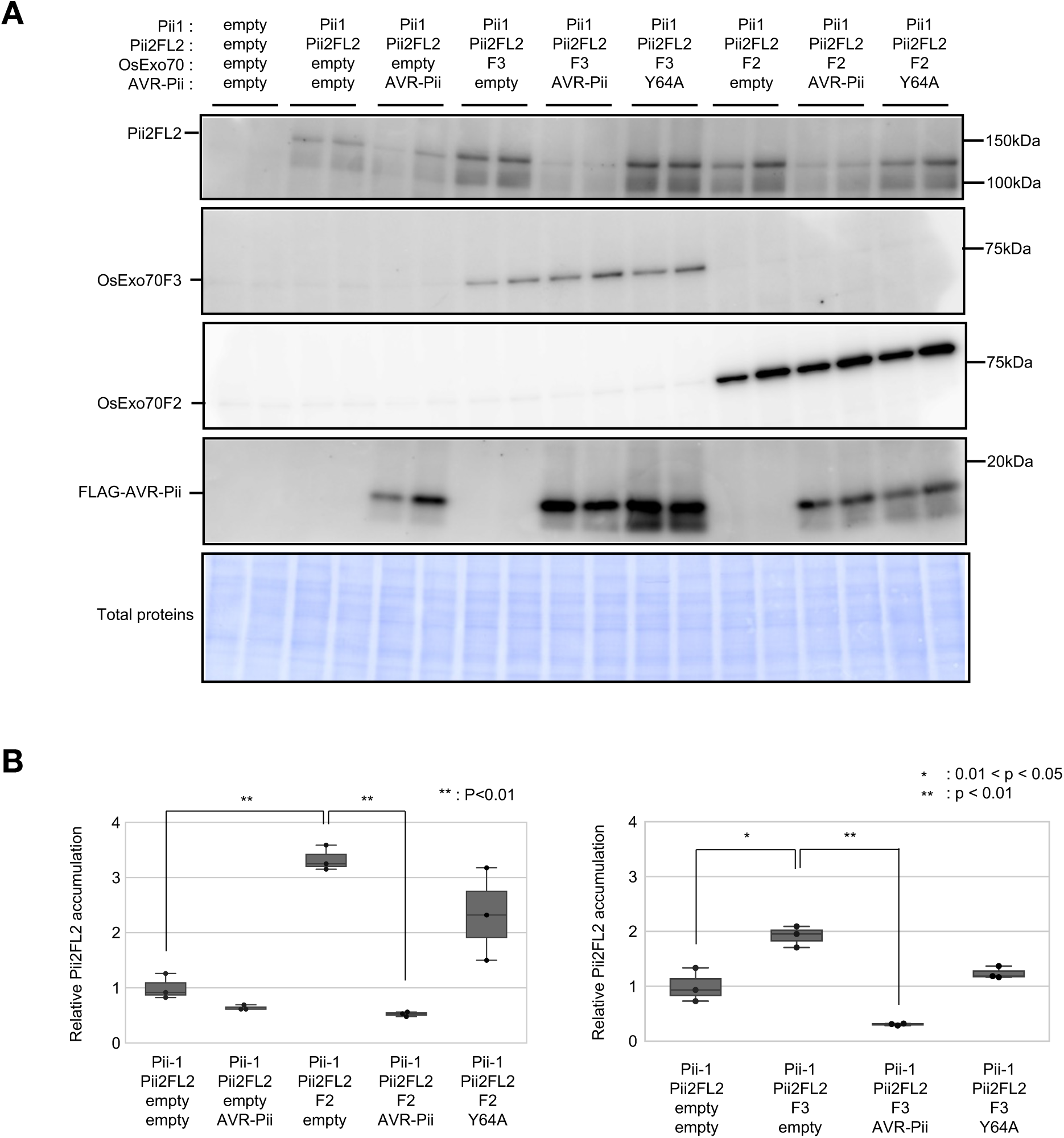
Effects of co-expression of OsExo70F2/F3 and AVR-Pii on Pii2FL2 accumulation. (A) Representative data of the accumulation of the indicated proteins in the transfected protoplasts prepared from rice Oc cell culture. AVR-Pii^Y64A^ (labeled Y64A: a loss-of-function mutant of AVR-Pii; see Fig. S7) was used as negative control. Pii2FL2 and FLAG-AVR-Pii were detected by immunoblot analysis using anti-FLAG antibodies. OsExo70F2 and OsExo70F3 were detected using a specific antiserum. The combinations of transfected constructs are indicated above. (B) Quantification of the abundance of Pii2FL2 encoded by the *Pii2FL2* construct, co-expressed with *AVR-Pii*, *AVR-Pii^Y64A^*, *OsExo70F2* (left), and/or *OsExo70F3* (right). Protein band intensity was quantified using ImageJ (https://imagej.net/ij/). Boxplots were illustrated from the values of three technical replicates. Single (*t*-test; 0.01<*P*<0.05) and double (*t*-test; *P*<0.01) asterisks represent significant differences of Pii2FL2 accumulation levels.

### AVR-Pii dissociation of the Pii2–OsExo70 complex triggers Pii2-mediated resistance

To elucidate the effect of AVR-Pii on the OsExo70–OsRIN4 and OsExo70–Pii2 interactions, we turned to an AlphaScreen. We first examined the effect of AVR-Pii on the OsExo70F2–OsRIN4c interaction. To this end, we first mixed FLAG-tagged OsRIN4c and biotinylated OsExo70F2 before adding AVR-Pii to study its effect on the preassembled OsRIN4c–OsExo70F2 complex (Fig. 5A). After AVR-Pii was added, the AlphaScreen signal intensity decreased in a dose-dependent manner (Fig. 5B, Fig. S21). This result suggests that AVR-Pii dissociates the OsRIN4c–OsExo70F2 complex, possibly representing the function of AVR-Pii as an effector. We performed a similar analysis using AVR-Pii and preassembled Pii2–OsExo70F2 and Pii2–OsExo70F3 complexes (Fig. 5A). After the addition of AVR-Pii, the AlphaScreen signal intensity decreased in a dose-dependent manner (Fig. 5C, Figs. S8), mirroring the AVR-Pii effect on the RIN4–OsExo70 interactions. In a series of AlphaScreens, AVR-Pii^Y64A^, which does not bind OsExo70, did not disrupt the interaction of OsExo70 with OsRIN4c or Pii2. From these results, we propose that AVR-Pii manipulates plant immunity *via* altering the RIN4–Exo70 interaction, and that Pii monitors this AVR-Pii action by presenting a NOI ID–Exo70 interaction decoy (Fig. 6).

**Figure 5.**
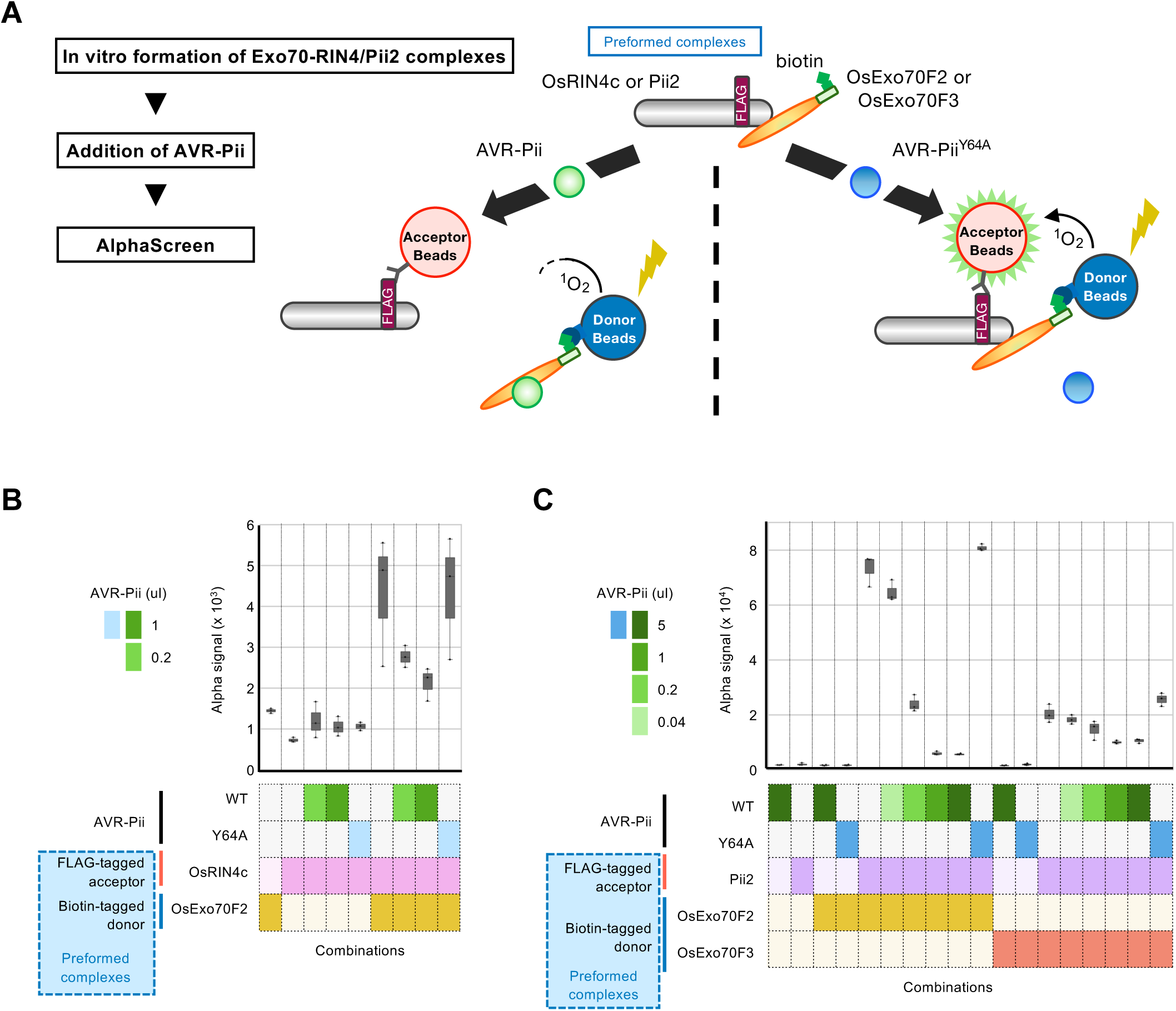
AVR-Pii dissociates the Pii2–OsExo70F2 and Pii2–OsExo70F3 complexes for *Pii2*-mediated resistance. (A) Scheme of AlphaScreen used to detect the effect of AVR-Pii on the interaction of OsRIN4c with OsExo70F2 and that of Pii2 with OsExo70F2. OsRIN4c and OsExo70F2 form a complex *in vitro*, as does Pii2 with OsExo70F2 (Fig. S6). Dissociation of the OsRIN4c–OsExo70F2 or Pii2–OsExo70F2 complex occurs upon addition of AVR-Pii, increasing the distance between the donor bead (bound to OsRIN4 or Pii2) and acceptor bead (bound to OsExo70F2) and thus decreasing the emitted AlphaScreen signals. As the AVR-Pii^Y64A^ mutant does not bind to OsExo70F2, the Pii2–OsExo70F2 complex remains intact, thus producing high AlphaScreen signals. The same assay was conducted with OsExo70F3 in place of OsExo70F2. (B) Effects of AVR-Pii on the preformed OsRIN4c– OsExo70F2 complex. FLAG-tagged OsRIN4c (FLAG-OsRIN4c) and biotinylated OsExo70F2 synthesized via a wheat germ extract (WGE) system were mixed to form the FLAG-OsRIN4c– OsExo70F2 complex (preformed complex), after which the indicated amount of AVR-Pii or AVR-Pii^Y64A^ produced from a WGE system was added to the mixture. After 1 h incubation at room temperature, the FLAG-OsRIN4c–OsExo70F2 interaction was detected by AlphaScreen. WGE without mRNA was used as negative control (indicated as clear rectangles). (C) Effects of AVR-Pii on the preformed Pii2–OsExo70F2 and Pii2–OsExo70F3 complexes. The indicated amount of AVR-Pii or AVR-Pii^Y64A^ produced from a WGE system was added to the preformed complex of Pii2FL2 and biotinylated OsExo70F2 or OsExo70F3, and the OsExo70F2–Pii2FL2 and OsExo70F3–Pii2FL2 interactions were detected by AlphaScreen as described above.

**Fig. 6.**
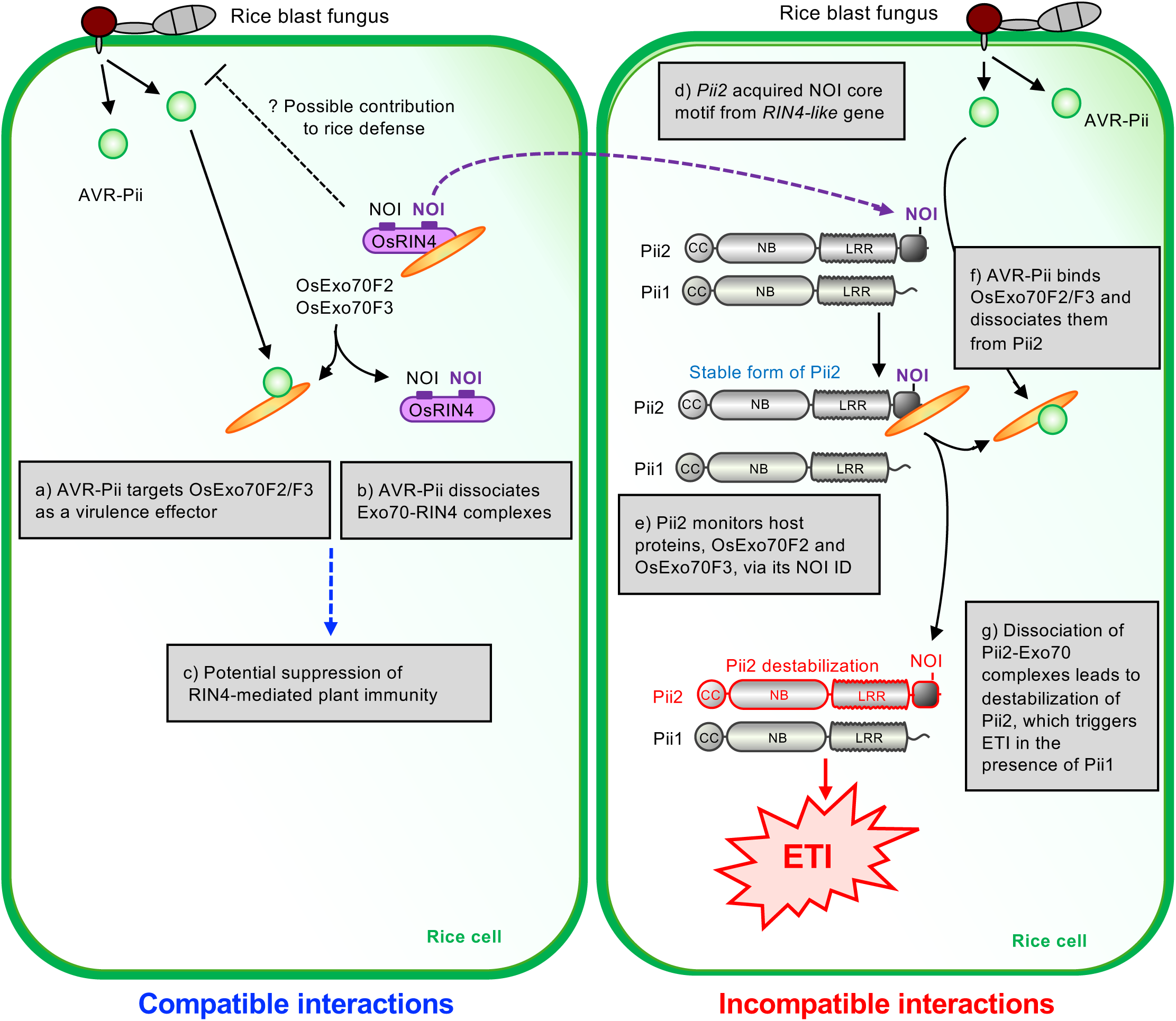
A possible model of Pii functions. In the compatible interaction between rice and *M. oryzae* (left panel), the fungal effector AVR-Pii is translocated to the cytoplasm of host cells during infection [Sharma et al., 2013]; and a) AVR-Pii binds host OsExo70F2 and OsExo70F3 [Fujisaki et al., 2015] tethered to OsRIN4-like proteins (which may potentially contribute to rice defense); b) the interactions between AVR-Pii and OsExo70F2/F3 results in dissociation of OsRIN4 from OsExo70F2/F3; c) this may lead to possible suppression of plant immunity; however, this should be confirmed in future study. In the incompatible interaction between rice and *M. oryzae* involving Pii (right panel); d) Pii2 evolutionarily acquired an NOI core motif (PxFGxW) as an integrated domain (ID) from the C-terminal PxFGxW motif of rice RIN4-like proteins; e) Pii2 binds OsExo70F2 and OsExo70F3 via its NOI core motif in the ID, which stabilizes Pii2; f) AVR-Pii binding to OsExo70F2/F3 leads to dissociation of OsExo70F2/F3 from Pii2; g) Dissociation of OsExo70F2/F3 from Pii2 destabilizes Pii2, resulting in the activation of effector-triggered immunity (ETI). The activation of immune responses may be cooperatively executed by Pii1 and Pii2 (Takagi et al., 2017).

## Discussion

The current view of plant immunity is that IDs of NLR immune receptors directly bind to or serve as substrates for pathogen effectors. Our findings suggest that NLR-IDs may detect host proteins targeted by pathogen effectors in a previously unrecognized mechanism whereby NLR-ID mediates indirect recognition of a pathogen effector [Dangl and Jones, 2001; van der Hoon and Kamoun, 2008] (Fig. 6). This model does not match any of the nine mechanisms of plant disease resistance proposed previously [Kourelis and van der Hoorn, 2018]. In *Arabidopsis*, RIN4 is a crucial regulator of plant immunity [Ray et al., 2019]. Considering the recent reports of RIN4–EXO70 interactions [Afzal et al., 2013; Sabol et al., 2017; Wu et al., 2021], we propose that rice RIN4–EXO70 complexes are the target of AVR-Pii as a virulence effector (Fig. 6). We hypothesize that 1) the Pii2– OsExo70F2 and Pii2–OsExo70F3 complexes are sensors and 2) the dissociation of Pii2 from OsExo70F2 or OsExo70F3 by AVR-Pii activates Pii2 (unstable form) to trigger immune responses. (Fig. 6). Based on the previous reports, the activation of immune response seems to be cooperatively executed by Pii1 and Pii2 (Takagi et al., 2013; Takagi et al., 2017). Current helper / sensor model for paired NLRs suggests that Pii2 functions as the sensor NLR, which may activate the helper NLR Pii1. The effects of altered Pii2 status on Pii1 function is the next important issue to understand the regulatory mechanism of the Pii NLR system.

Our work also points to the NOI–EXO70 complex as a major target of plant pathogen effectors. It is perhaps not surprising that these domains have been acquired as sensor domains by NLR immune receptors, with NOI and EXO70 commonly seen integrated into NLRs [Sarris et al., 2016; Brabham et al., 2017]. Whether integrated NOI and EXO70 domains can mediate the direct recognition of effectors remains to be determined. Nonetheless, our finding that NLR-IDs can indirectly detect AVR effectors expands the mechanistic view of pathogen detection by immune receptors: that is, the interactors of NLR-IDs can be either pathogen effectors or host proteins that are targeted by effectors. This work presents the first example to our knowledge of indirect pathogen recognition *via* the ID of an NLR, highlighting how the arms race between plants and pathogens has driven the emergence of a wealth of molecular interactions and mechanisms.

## Materials and Methods

### Plants and pathogens

*Pii2* and *OsExo70F2* knockout lines (Hit5882 and Hit3025, respectively) were isolated from EMS-treated mutant lines of the rice (*Oryza sativa*) cultivar Hitomebore [Takagi et al., 2013, 2017]. *OsExo70F3* knockout plants were generated using the CRISPR/Cas9 system with the pCRISPR/Cas9-F3-1 plasmid. *Agrobacterium tumefaciens* (EHA105) was transformed with the plasmid and used for stable transformation of Hitomebore. Transformation and regeneration of rice plants were performed according to Hiei et al. (1994). *Osexo70f2 Osexo70f3* double knockout lines were generated by pCRISPR/Cas9-F3-1-mediated knockout of *OsExo70F3* in the Hit3025 (*Osexo70f2* knockout) background.

To generate transgenic rice plants expressing *Pii2FL1*, *Pii2FL2*, *Pii2FL3*, or *Pii2FL2-m1*, the plasmids pCAMBIA-Pii2FL1, pCAMBIA-Pii2FL2, pCAMBIA-Pii2FL3, and pCAMBIA-Pii2FL2m1, respectively, were introduced to *A. tumefaciens* EHA105, which was subsequently used for transformation of Hit5882, a *pii2* mutant of the rice cultivar Hitomebore [Takagi et al., 2017].

*Magnapothe oryzae* isolate Sasa2 is stored at Iwate Biotechnology Research Center. A transgenic Sasa2 line expressing *AVR-Pii* (Sasa2-AVR-Pii) was previously established as Sasa2(+22p:pex33) [Yoshida et al., 2009]. For generating transgenic Sasa2 lines expressing *AVR-Pii* mutants (m9-9, Y64D, and Y64A), the plasmids pCB-pex22p-AVR-Pii^m9-9^, pCB-pex22p-AVR-Pii^Y64D^, and pCB-pex22p-AVR-Pii^Y64A^, respectively, were used for transformation of Sasa2 following the method described by Sugihara et al. (2023).

### Generation of AVR-Pii mutants

For generation of *AVR-Pii* mutants, random mutagenesis of pGAD-AVR-Pii using a Diversify^TM^ PCR Random Mutagenesis Kit (Takara) was performed using primers KF777f and KF778r [Fujisaki et al., 2015]. The resulting PCR products were digested with *Eco*RI and *Bam*HI and introduced into pGADT7 (Clontech, CA, USA) by utilizing *Eco*RI and *Bam*HI sites. pGADT7 vectors carrying *AVR-Pii* mutants including pGAD-pex22p-AVR-Pii-m9-9 (AVR-Pii M4; De la Concepcion et al., 2022) were cloned and used for Y2H assays in combination with pGBKT7-OsExo70F3 to verify the effects of AVR-Pii mutations on the interaction with OsExo70F3 [De la Concepcion et al., 2022].

### Expression of rice genes encoding NOI core motifs

The amino acid sequence of the *Arabidopsis* RIN4 protein (NP_189143.2) was used as the query for a BLAST search to identify protein-coding genes containing the NOI core motif (PxFGxW) in rice. To analyze gene expression, a conidial suspension (5 × 10^5^ conidia/ml) of Sasa2 containing 0.002% (v/v) Tween 20 was sprayed onto Hitomebore leaves and the leaves were harvested at 3 dpi. Hitomebore leaves treated with distilled water with 0.002% (v/v) Tween 20 (Mock) were used as a control. Total RNA of the harvested leaves was extracted using a NucleoSpin RNA Plant kit (Takara). cDNA was synthesized using the ReverTra Ace kit (TOYOBO, Osaka, Japan) with oligo dT primer. The primer sets used for RT-PCR of each gene are listed in Table S2. RT-PCR was performed as described previously [Fujisaki et al., 2015]. The rice actin gene (Os01g0866100) was amplified by RT-PCR and used as a control.

### Protein–protein interactions

Y2H assays were performed as described by Kanzaki et al. [2012]. Ten-times dilution series [OD_600_ = 3.0 (×1), 0.3 (×10^-1^), and 0.03 (×10^-2^)] of yeast cells were prepared and spotted onto quadruple dropout medium (QDO); basal medium lacking Trp, Leu, Ade, and His but containing 5-bromo-4-chloro-3-indolyl α-D-galactopyranoside (X-α-gal) (Clontech). To detect interactions, QDO medium both with and without 10 mM 3-amino-1,2,4-triazole (3AT) (Sigma) was used. Yeast cells were also spotted onto double dropout medium (DDO); basal medium lacking Trp and Leu to test cell viability. To determine protein accumulation in yeast cells, yeast cells were propagated in liquid DDO at 30°C overnight; 40 mg cells was collected by centrifugation at 15,000 × *g* for 10 sec, and resuspended with 160 μl GTN + DC buffer [10% (v/v) glycerol, 25 mM Tris-HCl (pH 7.5), 150 mM NaCl, 1 mM DTT, and 1 tablet of protease inhibitor (Complete EDTA-free; Roche, Basel Switzerland)) before adding 160 μl of 0.6 M NaOH, mixing gently, and incubating at room temperature for 10 min. Next, 160 μl of gel sample buffer [40% (w/v) glycerol, 240 mM Tris-HCl (pH 6.8), 8% (w/v) SDS, 0.04% (w/v) bromophenol blue, 400 mM DTT] was added and incubated at 95°C for 5 min. After centrifugation at 20,000 × *g* for 5 min, the supernatant was subjected to SDS-PAGE. Proteins expressed from bait and prey vectors were immunologically detected using anti-Myc-tag mAb-HRP-DirectT (MBL, Nagoya, Japan) and anti-HA-Peroxidase 3F10 (Roche), respectively. Co-IP experiments of transiently expressed proteins in *Nicotiana benthamiana* were performed as described by Fujisaki et al. [2015]. StrepII-HA (SH)-tagged OsExo70F2/F3 and FLAG-tagged OsRIN4c proteins were expressed using pGVG-SH-OsExo70F2, pGVG-SH-OsExo70F3, and pGVG-FL-OsRIN4c, respectively. At 18 h after agroinfiltration, 30 μM dexamethasone (DEX) was applied to infiltrated leaves to induce transgene expression. Leaves were harvested at 24 h after DEX treatment, and their homogenates in GTN buffer [Fujisaki et al., 2015] were used for co-IP. Anti-HA-Agarose affinity gel (Sigma) was used for co-IP of HA-tagged proteins, and proteins were eluted using 0.25 mg/ml HA peptide (Sigma).

For the AlphaScreen, *in vitro* transcription and protein synthesis were performed using a WEPRO7240 expression kit (Cell-Free Sciences, Kanagawa, Japan) according to the manufacturer’s instructions. cDNA fragments were amplified from several AlphaScreen plasmids carrying target genes using primers SPu and SP-A1868. *In vitro* transcripts were synthesized from each cDNA fragment using SP6 RNA polymerase in a 30-μl reaction mixture [6 μl of 5× transcription buffer (Cell-Free Sciences), 3 μl of 25 mM NTPs (Cell-Free Sciences), 0.3 μl of SP6 RNA polymerase (80U/μl; Cell-Free Sciences), 0.3 μl of RNase inhibitor (Cell-Free Sciences), 3 μl of cDNA fragment (direct use of PCR reaction mixture), 17.4 μl of distilled water]. After ethanol precipitation, transcripts were dissolved in 10 μl of distilled water and used for translation reactions performed in bilayer mode [Takai et al., 2010] for 16 h at 16°C.

To detect protein–protein interactions by AlphaScreen, *in vitro* assembly of protein complexes was carried out in a total volume of 15 μl containing 1 μl of biotinylated proteins, and 1 μl of FLAG-tagged proteins in the AlphaScreen buffer [100 mM Tris-HCl (pH 8.0), 150 mM NaCl, 0.01% (v/v) Tween 20, 1 mg/ml BSA] at 25°C for 1 h in a 384-well Optiplate (PerkinElmer). For competition assays, protein complexes were first assembled in a total volume of 10 μl containing 1 μl of biotinylated proteins (OsExo70F2/F3) and 1 μl of FLAG-tagged protein (Pii2FL2 or FL-OsRIN4c) in the AlphaScreen buffer (preformed complex). Competitor proteins (AVR-Pii and AVR-Pii^Y64A^) were diluted in a 1/5, 1/25, or 1/125 ratio with translation reaction mixture [Takai et al., 2010] without mRNA, and 5 μl of competitor solution containing 1 μl of a series of diluted competitor or undiluted competitor (corresponding to 0.008–1 μl of undiluted competitor) was prepared in AlphaScreen buffer. After incubation of the preformed complexes at 25°C for 1 h in a 384-well Optiplate, 5 μl of competitor solution was added and the resulting 15 μl of protein mixture was incubated at 25°C for 1 h in a 384-well Optiplate.

Following the instruction manual of the AlphaScreen IgG (Protein A) detection kit (PerkinElmer), 10 μl of detection mixture containing AlphaScreen buffer, 5 μg/ml anti-FLAG M2 antibody (Sigma-Aldrich), 0.08 μl of streptavidin-coated donor beads, and 0.08 μl of Protein A-coated acceptor beads were added to each well of the 384-well Optiplate, followed by incubation at 25°C for 1 hr. Luminescence was measured using the AlphaScreen detection program. Since OsRIN4c–OsExo70F2/F3 interactions were not detected by the AlphaScreen in the presence of 150 mM NaCl, OsRIN4c-mediated interactions were analyzed in the absence of NaCl. Mean values and standard deviations of each complex were calculated using the signals from three wells. All results were confirmed in two or more independent experiments.

### Assays for fungal pathogenicity and gene expression

Rice leaf blade spot inoculation with conidial suspension (5 × 10^5^ conidia/ml) was performed as described by Sugihara et al. [2023]. Disease lesions were photographed 10– 12 days post-inoculation (dpi), and their vertical length was measured. To evaluate the expression of transgenes in rice blast fungus carrying mutant AVR-Pii (Fig. S7), rice leaf tissues (cv. Moukoto) around the disease lesions at 10 days after spot inoculation were harvested, and RT-PCR was performed using primer set KF777f/KF778r. The *actin* gene (MGG_03982T0) of the blast fungus was detected by RT-PCR using primers MGactinF and MGactin R as a control.

### Transient gene expression in rice protoplasts

Rice protoplasts were isolated from rice cell culture (Oc cell) [Ishikawa et al., 2014; Ichimaru et al., 2022]. For cell wall digestion, 5 ml of packed cultured cells was mixed with 25 ml of cellulase solution [4% Cellulase RS (w/v; Yakult, Tokyo, Japan), 4% Cellulase R10 (w/v; Yakult), 0.1% (w/v) CaCl_2_.6H_2_O, 0.4 M mannitol, 0.1% (w/v) MES pH 5.6], and incubated with gentle shaking (60 rpm) at 30°C for 3 h. After filtration through Miracloth (Millipore, Billerica, MA), protoplasts in the flow-through fraction were collected by centrifugation at 800 × *g*, washed two times with 20 ml of W5 buffer (154 mM NaCl, 125 mM CaCl_2_, 5mM KCl, and 2mM MES pH 5.7), and adjusted to the desired concentration (2–3 × 10^6^ protoplasts/ml) with W5 buffer. For transfection, 10 μl of plasmid DNA containing 20 μg of the indicated combination of expression vectors (e.g., 4 μg of pAHC-FL-Pii1, 8 μg of pAHC-Pii2FL2, 2 μg of pAHC-OsExo70F2, 2 μg of pAHC-OsExo70F3, 2 μg of pAHC-AVR-Pii, and 2 μg of pAHC-luciferase) was mixed with 100 μl of protoplast solution. Next, 110 μl of PEG solution [40% PEG4000 (w/v; Fluka), 0.2 M mannitol, 0.1 M CaCl_2_] was added to the protoplast solution, mixed gently, and incubated at room temperature for 20 min. W5 buffer (700 μl) was then added, and protoplasts were collected by centrifugation at 800 × *g*. The protoplasts were washed with 700 μl of W5 buffer, resuspended in 500 μl of W5 buffer, and incubated in the dark at 30°C for 20 h. After incubation, protoplasts were collected by centrifugation at 800 × *g*, resuspended with 60 μl of 1 × GSB [62.5 mM Tris-HCl (pH6.8), 10% (v/v) glycerol, 0.2 g/ml SDS, 5μg/ml bromophenol blue, and 100 mM DTT], and subjected to immunoblotting. For cell death assays, the collected protoplasts were assayed using a Luciferase assay system (Promega, Madison, WI) according to the manufacturer’s instructions.

### Protein detection

Proteins were immunologically detected by immunoblot analysis. For detection of AVR-Pii protein without epitope tag, an anti-AVR-Pii antibody was produced using recombinant His-AVR-Pii-ns [Fujisaki et al., 2015] as antigen. Preparation of rabbit AVR-Pii-specific antiserum was performed by Scrum Inc. (Tokyo, Japan). OsExo70F2- and OsExo70F3-specific antisera [Fujisaki et al., 2015], respectively, were used to detect OsExo70F2 and OsExo70F3 without epitope tag expressed in rice protoplasts. Antibodies bound to the antigens were detected by HRP-conjugated anti-rabbit IgG (H+L) (Promega). HRP-conjugated anti-FLAG M2 (Sigma-Aldrich, St. Louis, MO), anti-HA 3F10 (Roche, Basel, Switzerland), and anti-Myc-tag mAb-HRP-DirectT (MBL, Tokyo, Japan) were used for the detection of FLAG-, SH-, and Myc-tagged proteins, respectively. Biotinylated proteins were detected using HRP-conjugated streptavidin (Cell Signaling, Danvers, MA).

### Phylogenetic analysis and BLAST search analysis

hmmsearch (https://www.ebi.ac.uk/Tools/hmmer/search/hmmsearch) was used against the “Reference Proteomes” for Poaceae (taxid: 4479) to search for NOI domain sequences. The profile HMM PF05627, known as the NOI domain and associated with the cleavage site for pathogenic type III effector avirulence factor Avr [Afzal et al., 2013], was used as the query profile. The following cut-offs were applied in the hmmsearch: -T 3 --domT 3 --incT 5 --incdomT 5. After obtaining the sequences containing NOI domains, those having any missing residues (‘X’) or more than two NOI domains were filtered out. The six validated rice NOI protein sequences were added to the 607 retained sequences, and the command-line hmmsearch v3.3.2 was executed again with the same cut-offs to generate a multiple sequence alignment of NOI domains in Stockholm format. This alignment was converted to FASTA format using esl-reformat. Sequences containing NOI motifs without any gaps (‘-’) were retained for phylogenetic analysis. IQ-TREE v2.2.0.3 [Minh et al., 2020; http://www.iqtree.org] with 1,000 ultrafast bootstrap replicates [Hoang et al., 2018] was used for phylogenetic analysis. The best-fit model to reconstruct the tree was automatically selected by ModelFinder [Kalyaanamoorthy et al., 2017] in IQ-TREE, which selected “Q.plant+I+G4” according to the Bayesian Information Criterion (BIC). Finally, NLR proteins were annotated using NLRtracker [Kourelis et al., 2021], and the resulting phylogenetic tree was visualized and annotated using iTOL [Letunic & Bork, 2021; https://itol.embl.de].

## Acknowledgements and funding sources

We thank Dr. John J.A. Dominguez, Dr. Hiroyuki Kanzaki, and Sayaka Fujisaki for technical and general assistance. This work was supported by JSPS Grants 15H05779, 20H05681, 23K20042 and 24H00010 to RT and 26850029 to KF. Biotechnology and Biological Sciences Research Council (BBSRC, UK) grants BB/X010996/1 to MJB and SK and V015508/1 to MJB.

## Footnotes

The authors declare no competing interests.

## Data, Materials, and Methods

All data presented in this study are included in the article and/or the *SI Appendix*. The datasets used for phylogenetic analysis were uploaded to Github (https://github.com/YuSugihara/NOI_domains) and Zenodo (https://doi.org/10.5281/zenodo.11216127). Detailed information on the construction of plasmids used in this study is provided in Supplementary text S1 and Table S1, and the primer sequences used for the experiments are listed in Table S2.

---

**Figure S1.**
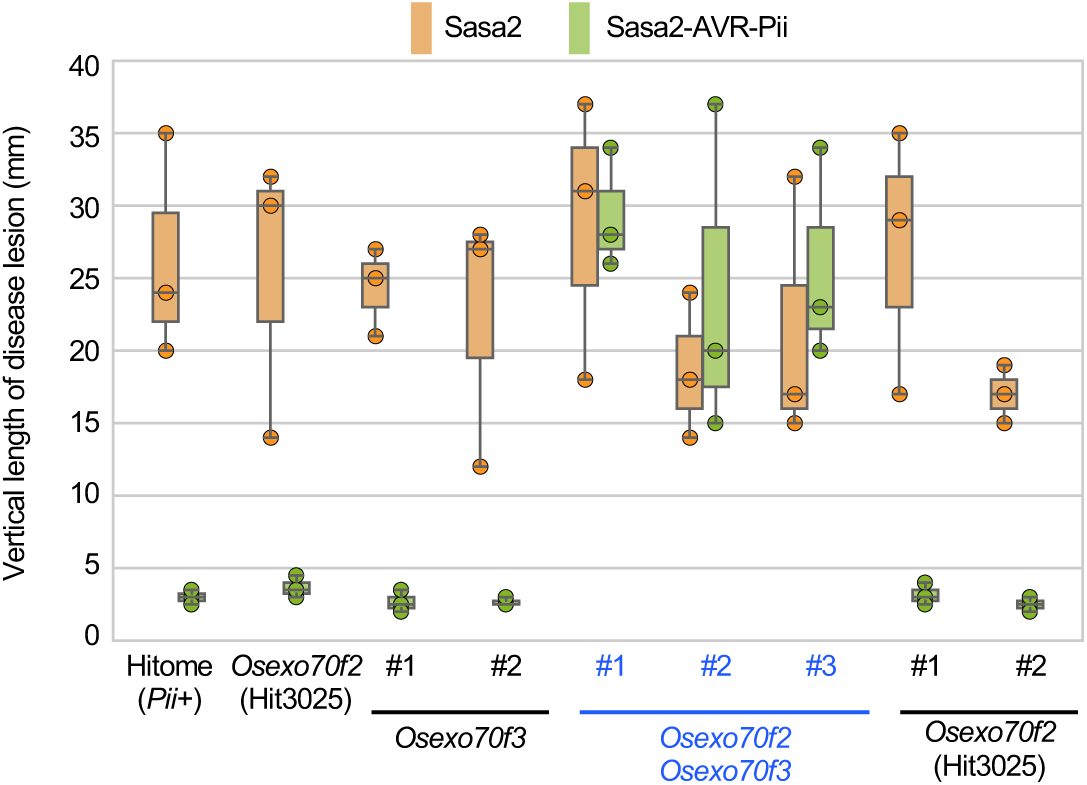
Quantified disease lesion length in rice lines with loss of OsExo70F2 and/or OsExo70F3 function after inoculation with *Magnaporthe oryzae* isolate Sasa2 or Sasa2-AVR-Pii. Rice blast fungus isolate Sasa2 (without *AVR-Pii*) and transgenic strain Sasa2–AVR-Pii [Yoshida et al., 2009] were used to inoculate rice plants of the single and double knockout mutant lines *Osexo70f2* and *Osexo70f3* in the Hitomebore background (described in Fig. 1). The vertical lengths of disease lesions were measured at 10 days post inoculation. Boxplots were illustrated from the values of three inoculated spots per line. In the *Osexo70f2 Osexo70f3*double knockout line (labeled in blue), *Pii*-mediated resistance against Sasa2-AVR-Pii was abrogated.

**Figure S2.**
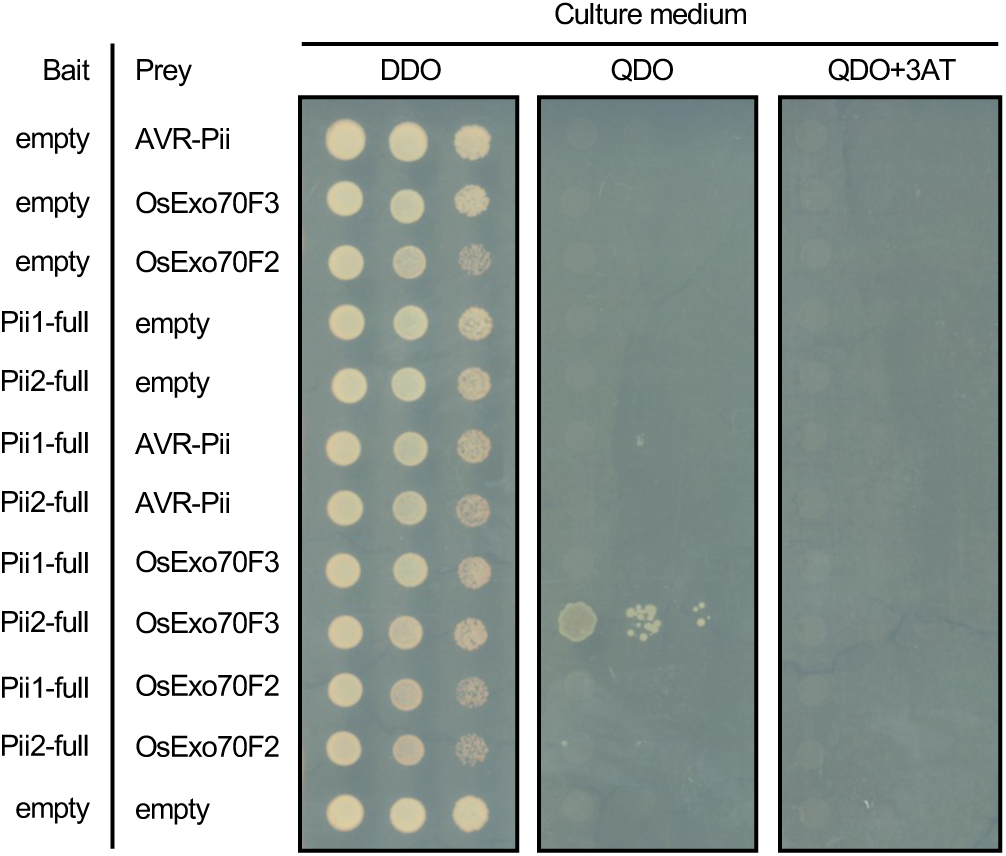
Yeast two-hybrid assay testing the interactions of Pii1 or Pii2 with OsExo70F2, OsExo70F3, or AVR-Pii. Ten-fold serial dilution of positive yeast transformants were spotted onto double dropout (DDO) medium as control and on quadruple dropout (QDO) medium and QDO medium + 10 mM 3-amino-1,2,4-triazole (3AT) to test interactions. Empty vector (empty) was used as negative control.

**Figure S3.**
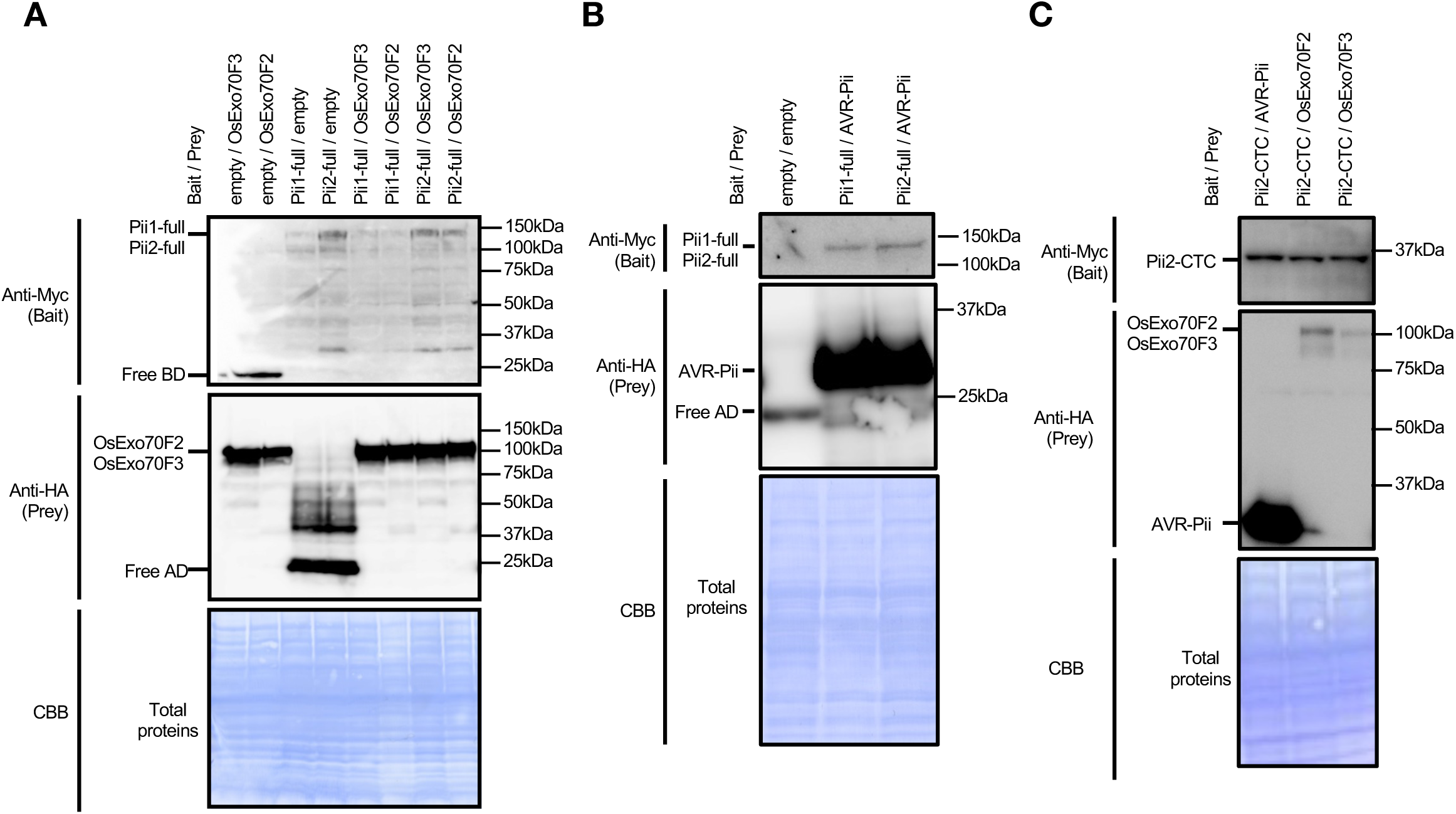
Confirmation of protein production for Pii1, Pii2, AVR-Pii, OsExo70F2, and OsExo70F3 in yeast two-hybrid assays. (A) Immunoblot analysis showing the accumulation of proteins with the expected molecular weight in yeast cells used in the yeast two-hybrid (Y2H) assays testing the interaction between full-length Pii1 or Pii2 and OsExo70F2 or OsExo70F3 (Fig. S2). The presence of proteins encoded by the bait and prey vectors was probed using anti-Myc and anti-HA antibodies, respectively, as shown in the upper (bait) and lower panels (prey). The expected position of proteins is indicated at left. A Coomassie brilliant blue (CBB)-stained proteins are shown below as loading control. (B) Immunoblot analysis showing the accumulation of proteins with the expected molecular weight in yeast cells used in the Y2H assay testing the interaction between AVR-Pii and full-length Pii1 or Pii2 (Fig. S2). (C) Immunoblot analysis showing the accumulation of proteins with the expected molecular weight in yeast cells used in the Y2H assay testing the interaction between AVR-Pii, OsExo70F2, OsExo70F3, and Pii-2-CTC (Fig. 2B).

**Figure S4.**
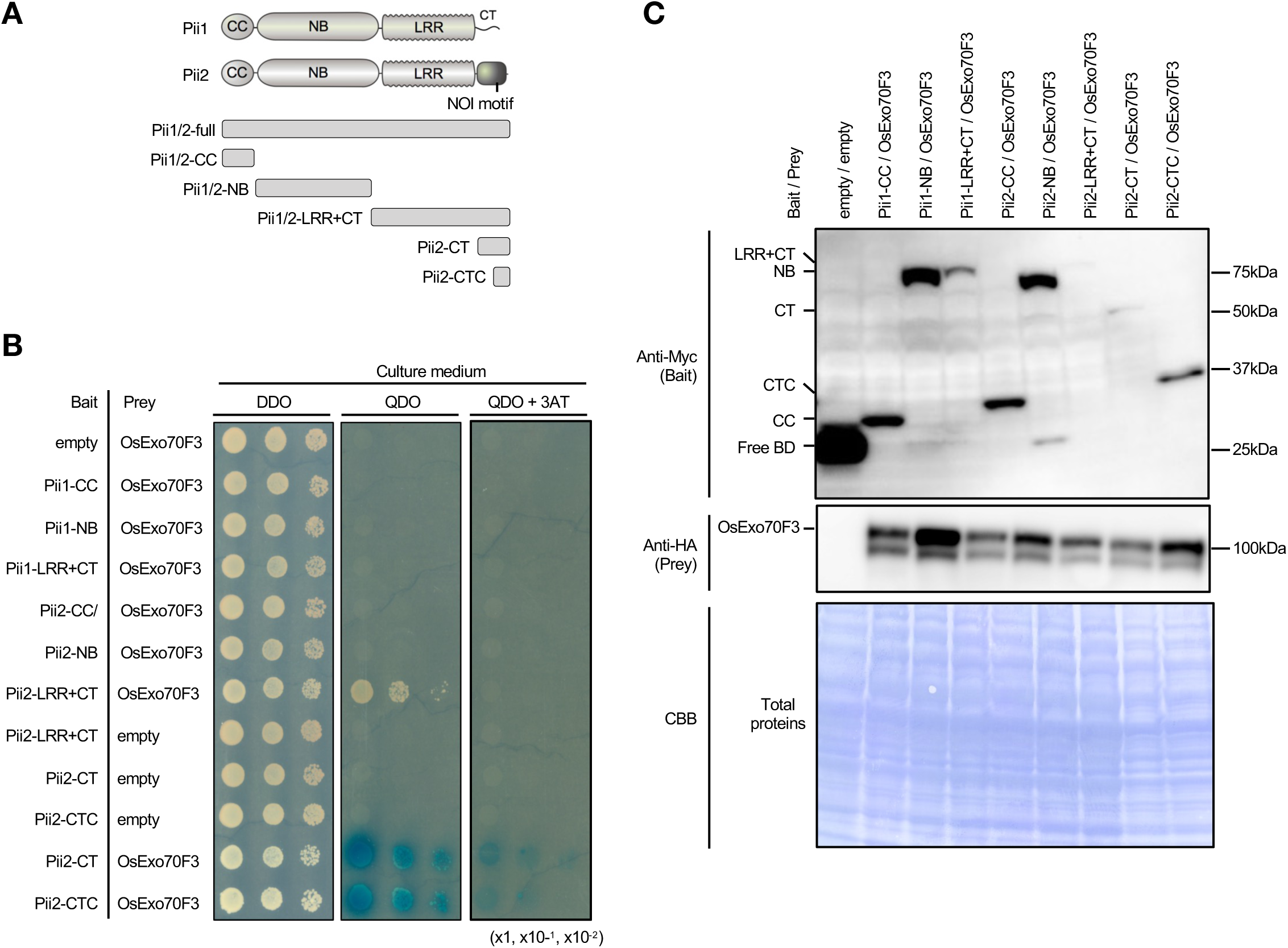
Mapping of the OsExo70F3-interacting regions of Pii2. (A) Diagram of the functional domains in Pii1 and Pii2 and their truncated variants used for mapping the OsExo70F3-interacting domain. (B) Y2H assay testing the interaction of OsExo70F3 with different regions of Pii1 and Pii2. Ten-fold serial dilution of positive yeast transformants were spotted onto DDO medium as control and on QDO medium and QDO medium + 10 mM 3AT to test interactions. Empty vector (empty) was used as negative control. Bait and prey combinations are listed at left. (C) Immunoblot analysis showing the accumulation of proteins with the expected molecular weight for Pii truncations and OsExo70F3 in yeast cells used in the Y2H assay. The presence of proteins encoded by the bait and prey vectors was probed using anti-Myc and anti-HA antibodies, respectively, as shown in the upper (bait) and lower panels (prey). The expected position of proteins is indicated on the left. A CBB-stained proteins are shown below as loading control.

**Figure S5.**
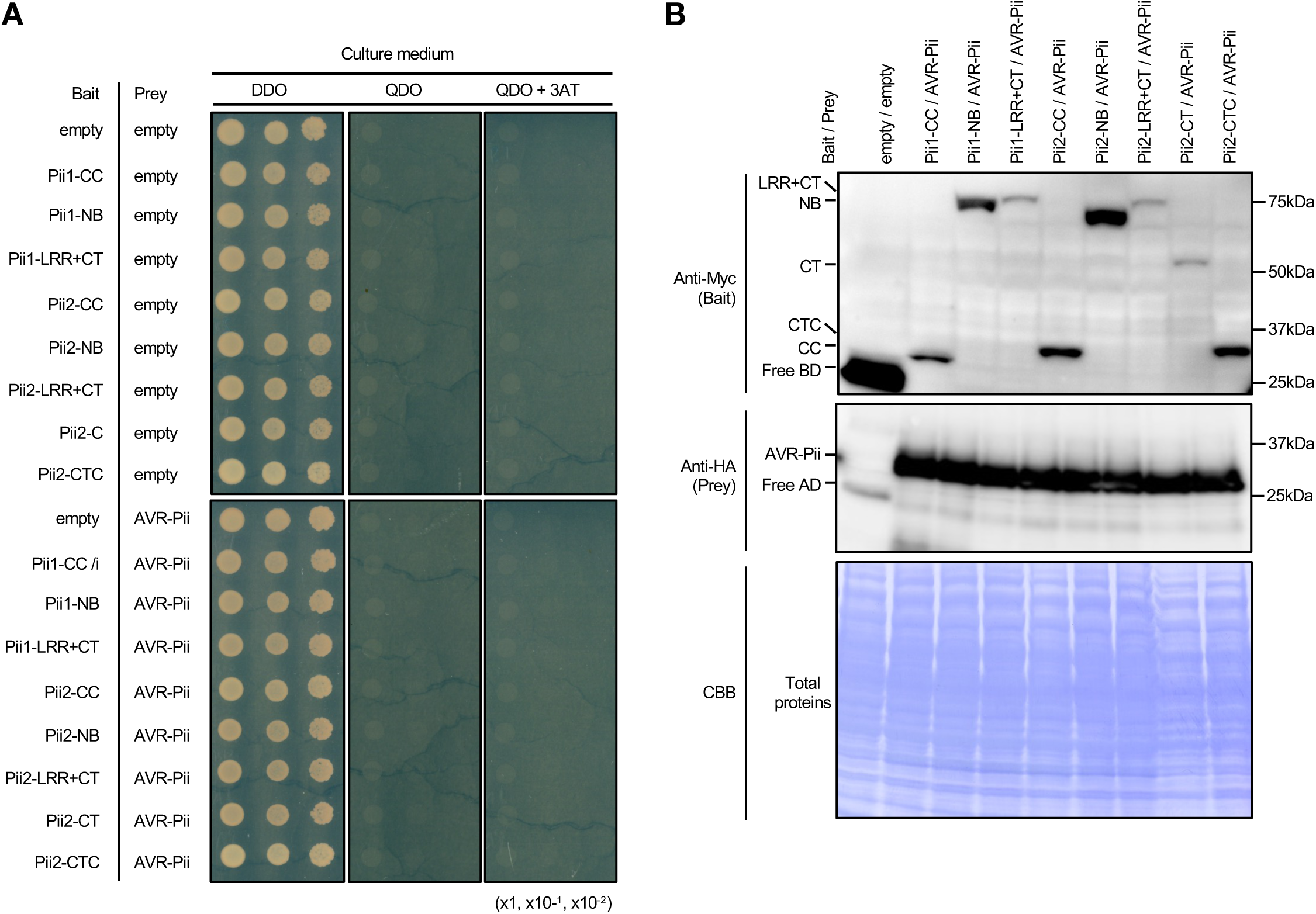
Interactions of AVR-Pii with Pii1 and Pii2 sub-fragments. (A) Y2H assay addressing the interactions of AVR-Pii with different regions of Pii1 and Pii2. Pii1 and Pii2 sub-fragments used in the assay are indicated in Fig. S4A. Ten-fold serial dilution of positive yeast transformants were spotted onto DDO medium as control and on QDO medium and QDO medium + 10 mM 3AT to test interactions. Empty vector (empty) was used as negative control. Bait and prey combinations are listed at left. (B) Immunoblot analysis showing the accumulation of proteins with the expected molecular weight for Pii truncations and AVR-Pii in yeast cells used in Y2H assay. The presence of proteins encoded by the bait and prey vectors was probed using anti-Myc and anti-HA antibodies, respectively, as shown in the upper (bait) and lower panels (prey). The expected position of proteins is indicated at left. A CBB-stained proteins are shown as loading control.

**Figure S6.**
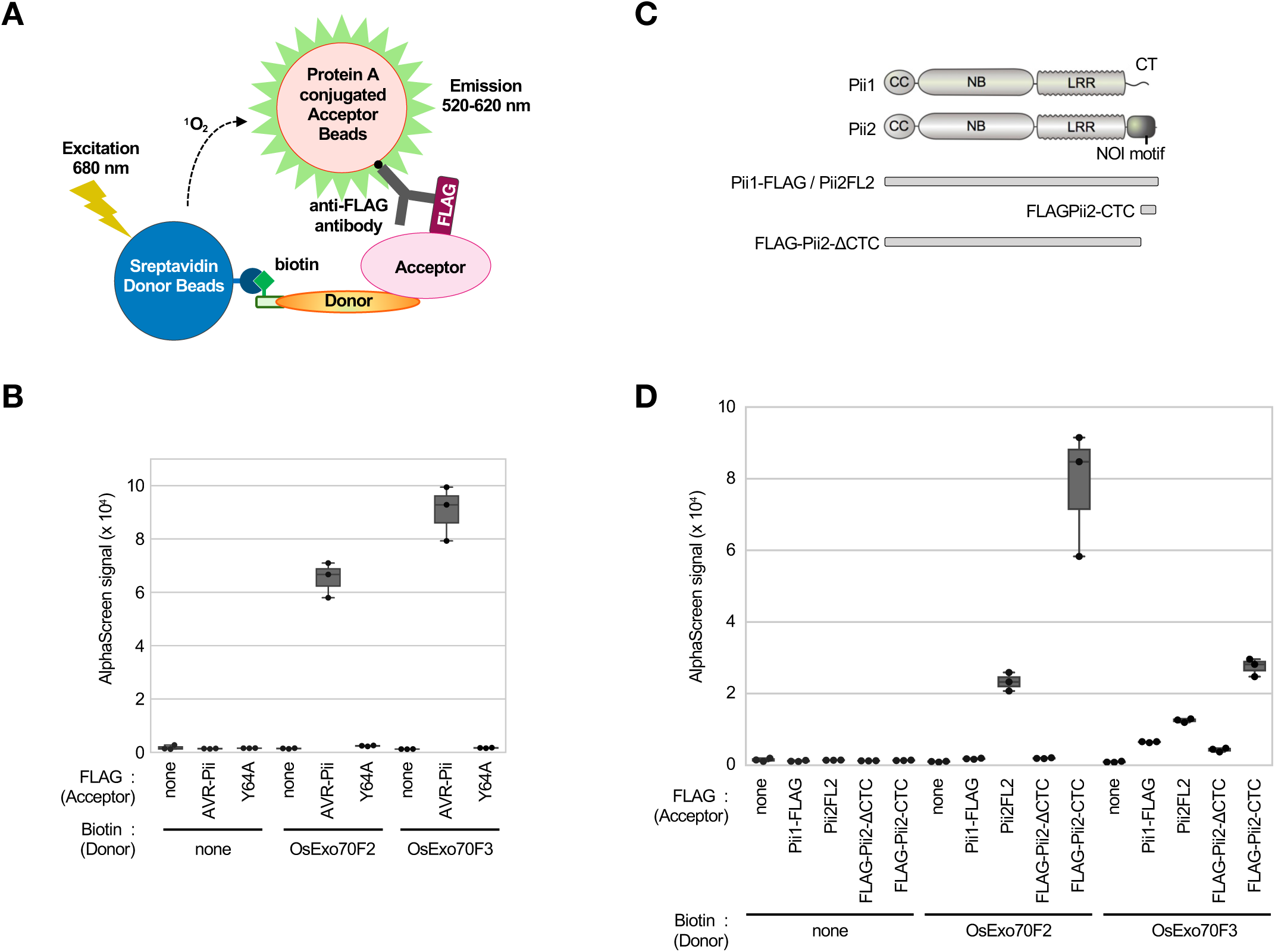
AlphaScreen to investigate the interactions of the components of the Pii system (Pii1, Pii2, OsExo70F2/F3, and AVR-Pii) (A) Principle of AlphaScreen. Donor beads are excited by illumination at 680 nm, leading to the conversion of ambient oxygen to singlet oxygen (^1^O_2_). Biotinylated proteins bind to streptavidin coating the donor beads, and protein A-conjugated acceptor beads bind to FLAG-tagged proteins via an anti-FLAG antibody. When biotinylated proteins and FLAG-tagged proteins form a complex, they bring the donor and acceptor beads in close proximity, allowing the singlet oxygen to be transferred to the acceptor beads, which become activated and emit fluorescence at 520–620 nm. (B) Interaction tests between biotinylated OsExo70F2 or OsExo70F3 and FLAG-tagged AVR-Pii by AlphaScreen. The AVR-Pii^Y64A^ mutant (Y64A), which does not interact with OsExo70F3 in a Y2H assay nor induces *Pii*-mediated resistance (Fig. S7), was used as negative control. WGE without mRNA added (none) was also used as negative control. Boxplots were illustrated from the values of three technical replicates. (C) Diagrams of full-length Pii1 and Pii2 showing their functional domains and the fragments used for AlphaScreen. (D) AlphaScreen to test the interaction between biotinylated OsExo70F2 or OsExo70F3 and FLAG-tagged full-length Pii1, Pii2, or Pii2 fragments (CTC and ΔCTC). All biotinylated and FLAG-tagged proteins were synthesized in wheat germ extracts (WGE) (Fig. S8), and the indicated WGE products were mixed in the indicated combinations for AlphaScreen. WGE without mRNA added (none) was used as negative control. Boxplots were illustrated from the values of three technical replicates.

**Figure S7.**
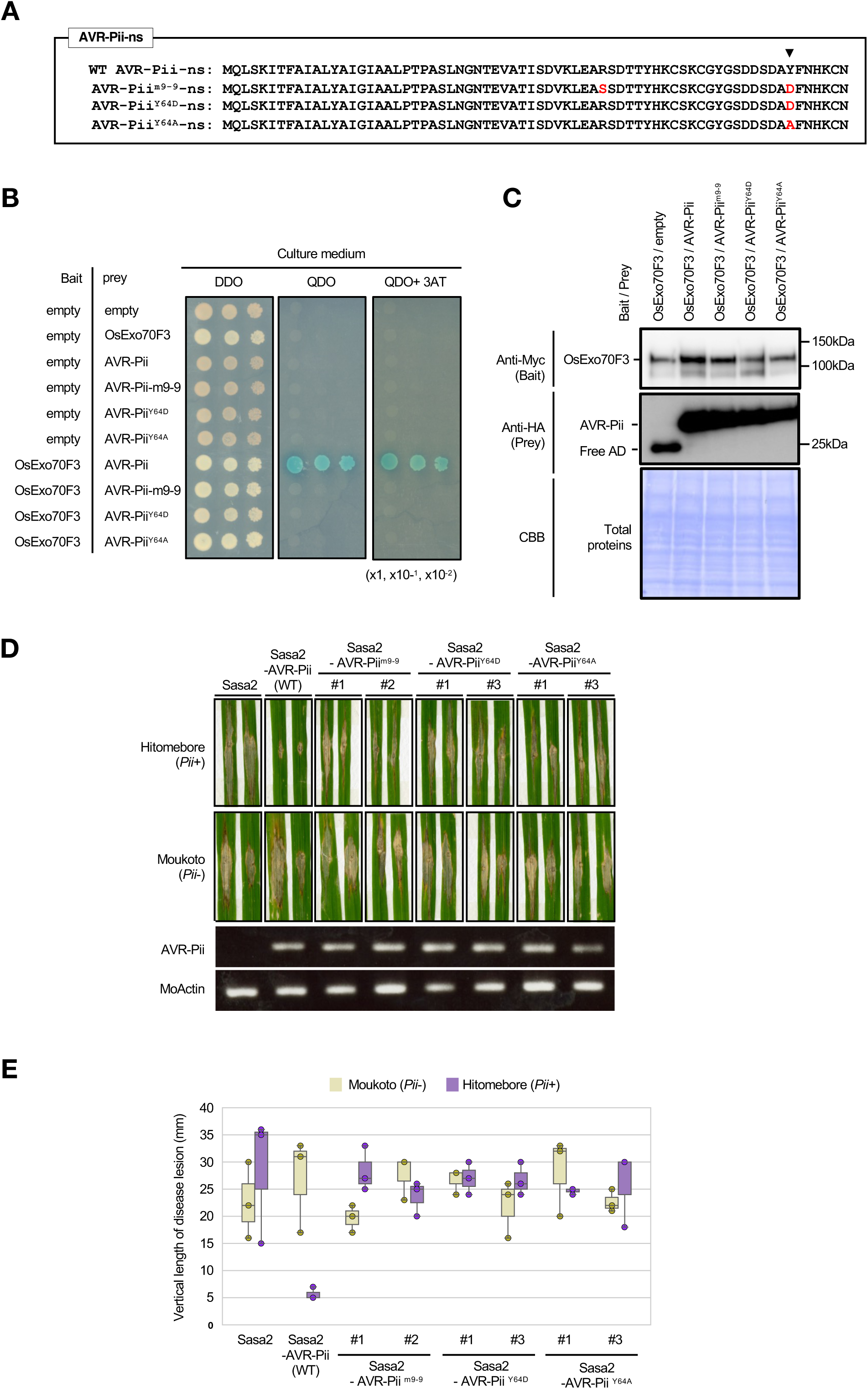
The AVR-Pii mutant AVR-Pii^Y64A^ does not bind OsExo70F3 and is not recognized by Pii. (A) Deduced amino acid sequence of mature AVR-Pii without signal peptide (AVR-Pii-ns) and related point mutants. The sequence of wild-type (WT) AVR-Pii-ns is given in the top row. Arrow head indicate the position of Tyrosine 64. The positions of mutations (m9-9, Y64D and Y64A) in AVR-Pii-ns are marked in red. The mutations were introduced with a Diversify^TM^ PCR random mutagenesis kit (Clontech); the resulting AVR-Pii mutants were used in the Y2H assay below. (B) Y2H assay showing the interaction between OsExo70F3 and wild-type AVR-Pii-ns, but not the AVR-Pii-ns mutants. Ten-fold serial dilution of positive yeast transformants were spotted onto DDO medium as control and on QDO medium and QDO medium + 10 mM 3AT to test interactions. Empty vector (empty) was used as negative control. Bait and prey combinations are indicated at left. (C) Immunoblot analysis showing the accumulation of proteins with the expected molecular weight for OsExo70F3 and wild-type and mutant versions of AVR-Pii-ns in yeast cells used in the Y2H assay. The presence of proteins encoded by the bait and prey vectors was probed using anti-Myc and anti-HA antibodies, respectively, as shown in the upper (bait) and lower panels (prey). The expected position of proteins is indicated on the left. A CBB-stained proteins are shown below as loading control. (D) AVR-Pii mutants (m9-9, Y64D, and Y64A) fail to induce Pii-mediated resistance. *M. oryzae* isolate Sasa2 (without *AVR-Pii*) and the transgenic strain Sasa2 expressing wild-type *AVR-Pii* (Sasa2–AVR-Pii; Yoshida et al., 2009) or AVR-Pii mutants (m9-9, Y64D and Y64A) were spot inoculated onto the rice leaves of the Moukoto (*Pii*−) and Hitomebore (*Pii*+) cultivars. Representative data of disease lesions at 10 days post inoculation are shown. Expression of *AVR-Pii* and *MoActin* was assessed by RT-PCR (bottom). (E) Boxplots showing disease lesion lengths after inoculation of the rice cultivars Moukoto and Hitomebore with *M. oryzae*, which is illustrated from the values of three inoculated spots per line. We used only AVR-Pii^Y64A^ for further study.

**Figure S8.**
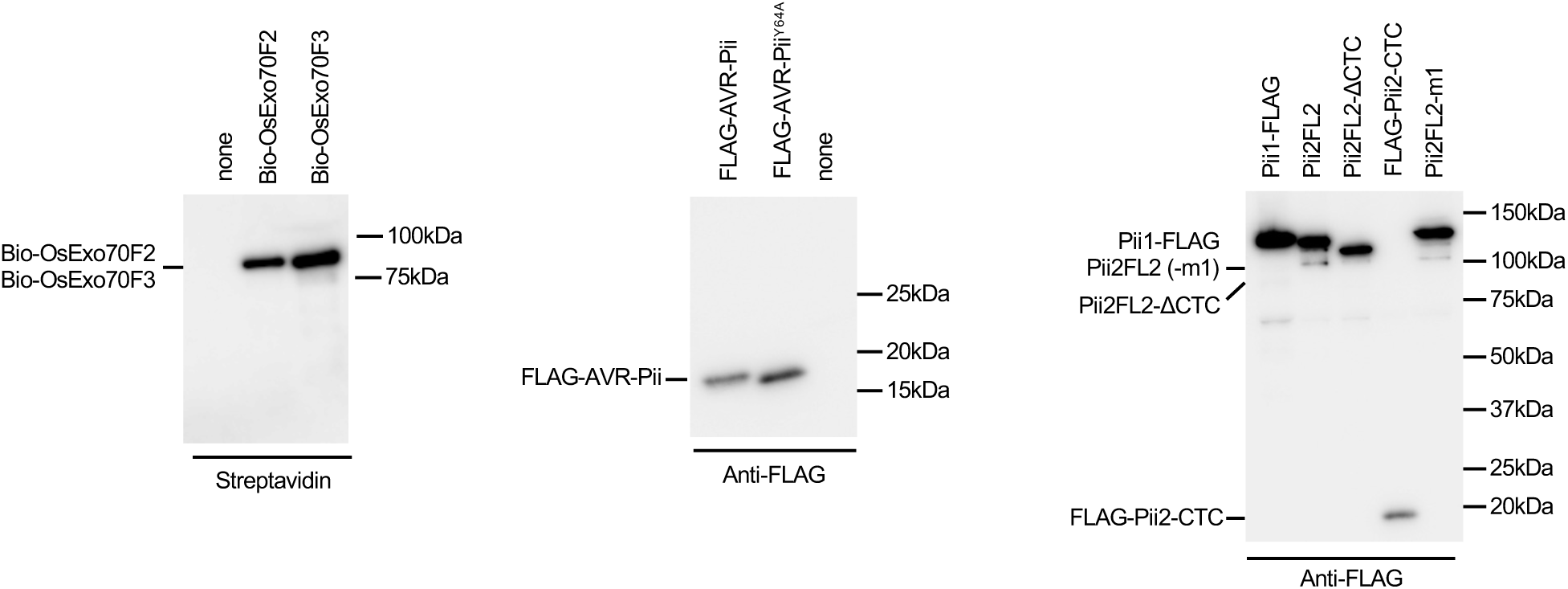
Confirmation of proteins synthesized in the wheat germ translation system used for AlphaScreen. Biotinylated OsExo70F2 or OsExo70F3 and FLAG-tagged AVR-Pii (FLAG-AVR-Pii), AVR-Pii^Y64A^ (FLAG-AVR-Pii^Y64A^), Pii1 (Pii1-FLAG), Pii2 (Pii2FL2), and Pii2 derivatives (Pii2FL2-ΔCTC, FLAG-Pii2-CTC and Pii2FL2-m1) were used for AlphaScreen in this study. Biotinylated (left) and FLAG-tagged (middle and right) proteins were detected using streptavidin and anti-FLAG antibodies, respectively.

**Figure S9.**
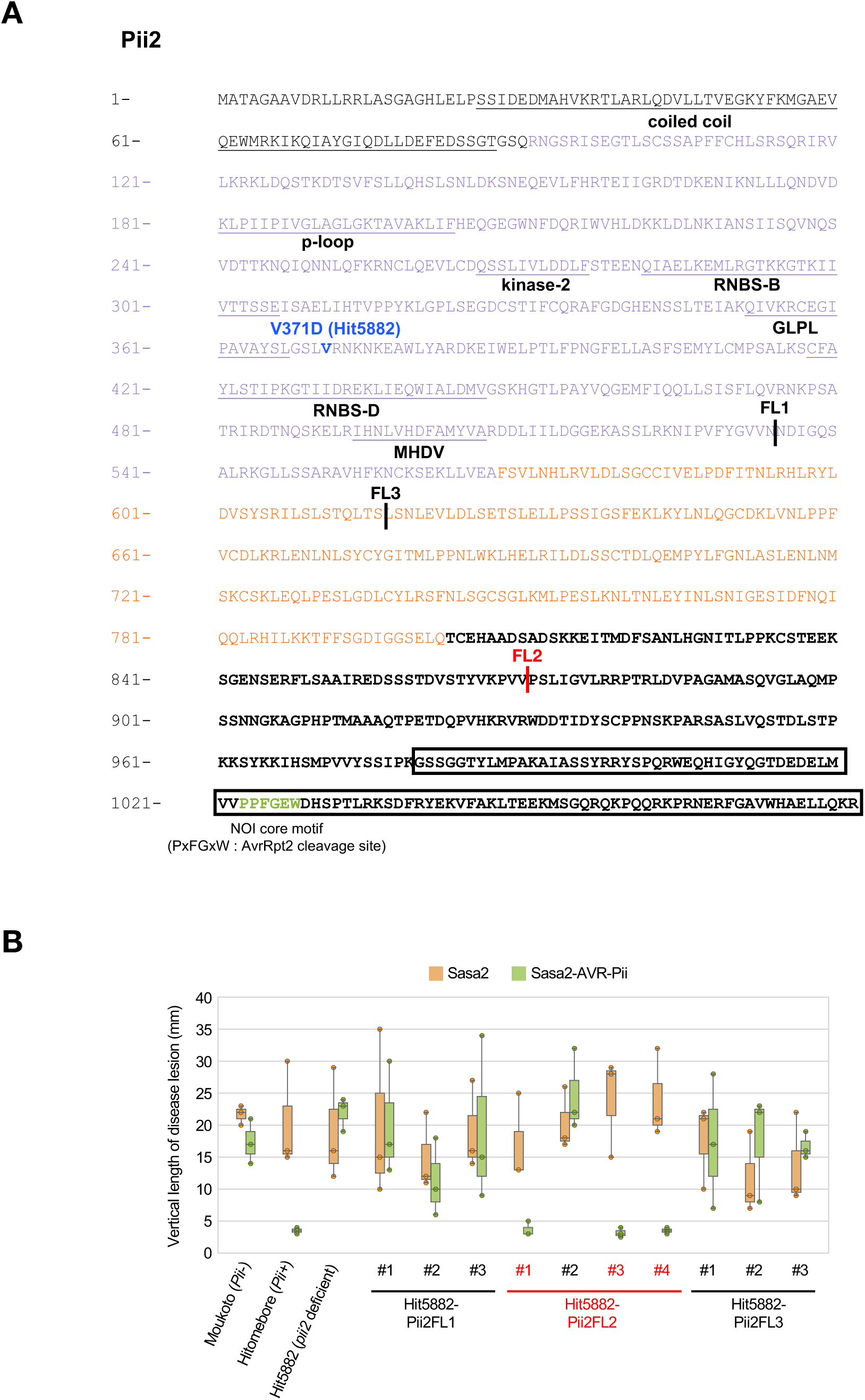
Amino acid sequence and domain structure of Pii2 and the effect of FLAG tag insertions on its AVR-Pii recognition function. (A) Deduced amino acid sequence of Pii2 from the rice cultivar Hitomebore. The NB and LRR domains are indicated as purple and orange letters, respectively. Coiled coil, P-loop, and other motifs described in Pi5-2 [Lee et al., 2009] are underlined. The NOI core motif (PxFGxW: AvrRpt2 cleavage site) is indicated by green letters. The amino acid residue corresponding to the V371D mutation in a *pii2* mutant (Hit5882) is marked in blue. The Pii2-CT region is indicated in black bold text and Pii2-CTC is indicated by a box. The three insertion sites for the FLAG-tag (FL1, FL2, and FL3) are also shown. (B) Lengths of disease lesions after spot inoculation of rice leaf blades with *M. oryzae* Sasa2 or Sasa2–AVR-Pii. The lesion lengths of Sasa2- and Sasa2–AVR-Pii-inoculated spots in the transgenic rice line Hit5882 expressing FLAG-tagged Pii2 inserted at different positions (*Pii2FL1*, *Pii2FL2*, and *Pii2FL3*) are shown. Boxplots were illustrated from the values of three inoculated spots per line. None of the lines of Hit5882-*Pii2FL1* or Hit5882-*Pii2FL3* showed resistance, whereas three lines of Hit5882-Pii2FL2 showed resistance. Therefore, the *Pii2FL2* construct was used in this study to retain Pii2 function in a FLAG-tagged form.

**Figure S10.**
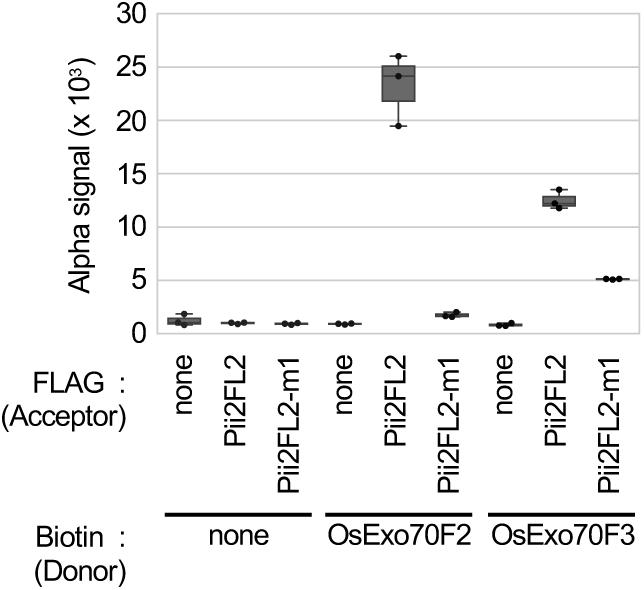
The NOI core motif of Pii2 ID is crucial for its interaction with OsExo70F2 and OsExo70F3. Interaction test between the Pii2-m1 mutant and OsExo70F2 or OsExo70F3 as studied by AlphaScreen. Boxplots were illustrated from the values of three technical replicates. For the confirmation of proteins used in the assay, see Figure S8.

**Figure S11.**
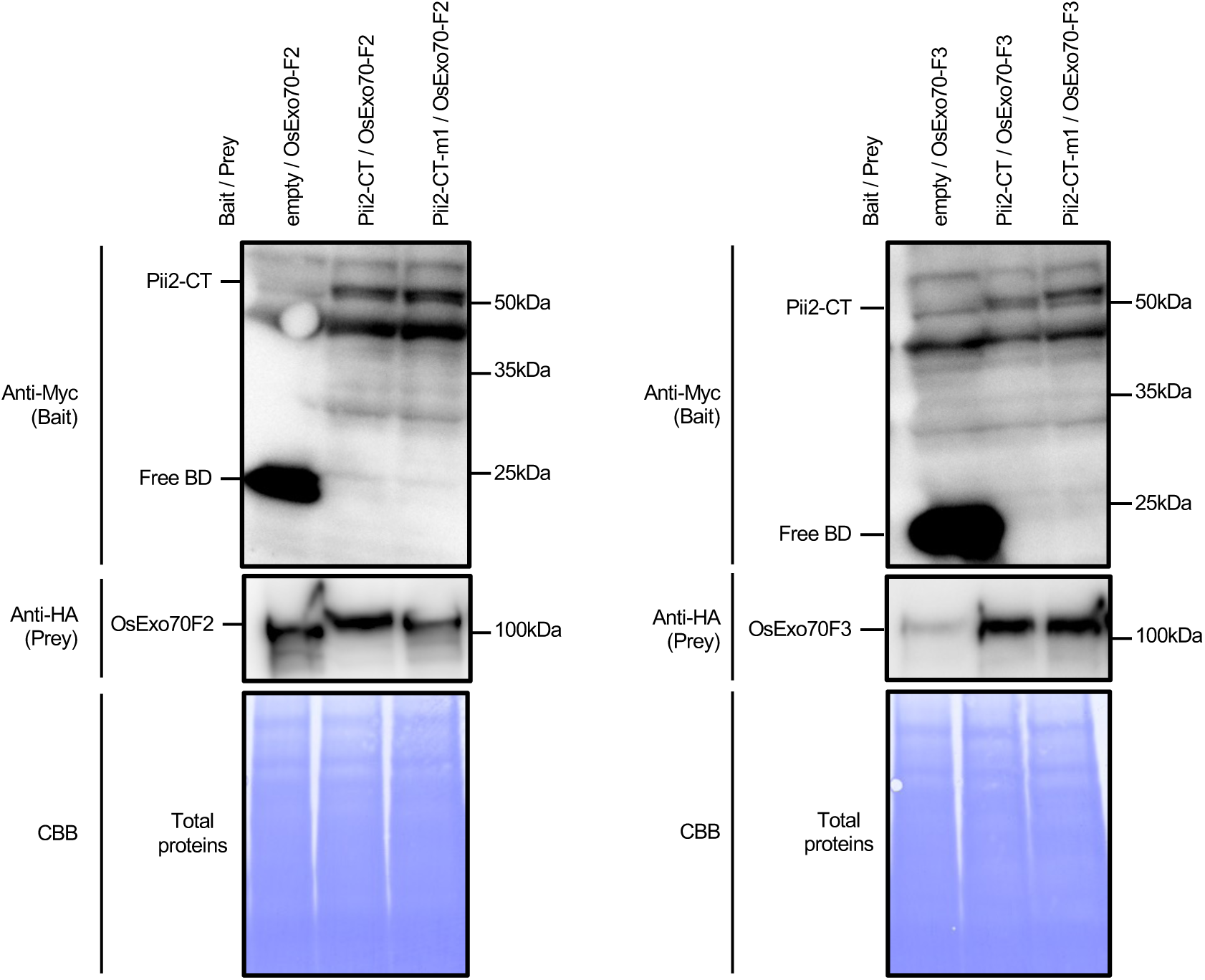
Confirmation of accumulation of OsExo70F2, OsExo70F3, Pii2-CT, and Pii2-CT-m1 proteins in yeast cells used for Y2H. Immunoblot analysis showing the accumulation of proteins with the expected molecular weight in yeast cells used in the Y2H assay (Fig. S10) testing the interaction between OsExo70F2 and Pii2-CT or Pii2-CT-m1 (left) or between OsExo70F3 and Pii2-CT or Pii2-CT-m1 (right). The presence of proteins encoded by the bait and prey vectors was probed using anti-Myc and anti-HA antibodies, respectively, and shown in the upper (bait) and lower panels (prey). The expected position of proteins is indicated on the left. A CBB-stained proteins are shown below as loading control.

**Figure S12.**
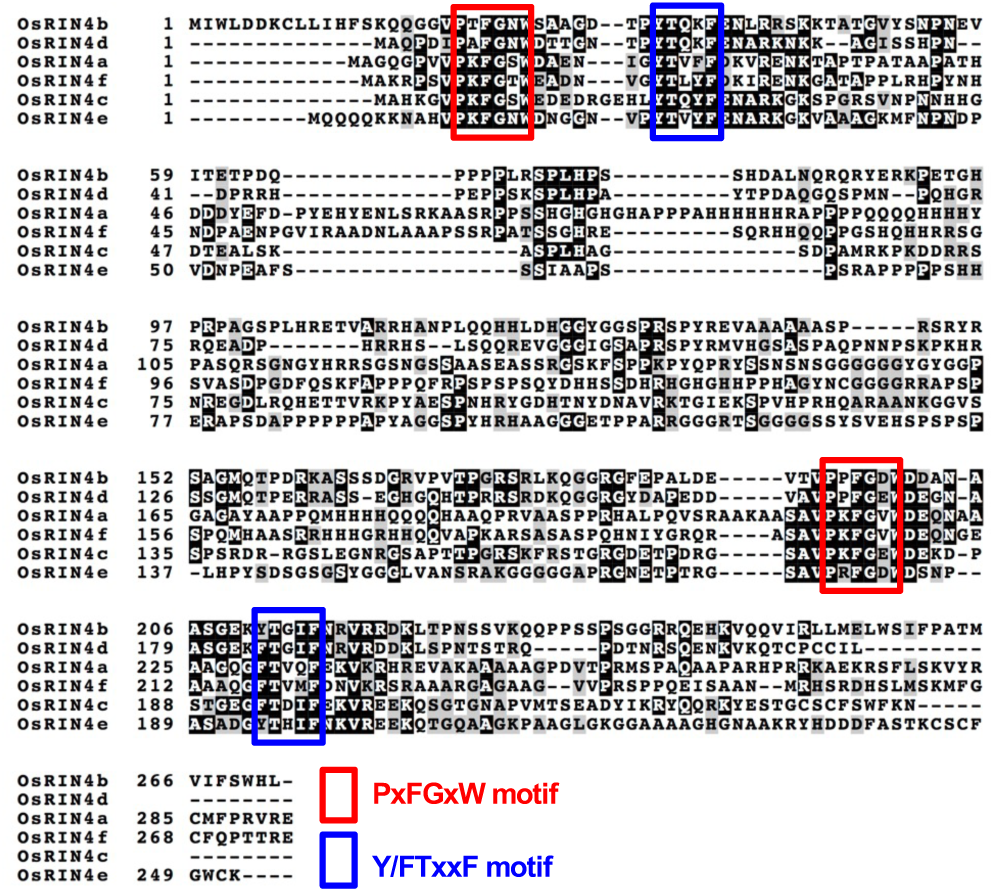
Multiple amino acid sequence alignment of rice RIN4-like proteins. Rice RIN4-like proteins, OsRIN4a (Os02g0189400), OsRIN4b (Os02g0504700), OsRIN4c (Os03g0848600), OsRIN4d (Os04g0379600), OsRIN4e (Os08g0526400), and OsRIN4f (Os06g0636100) were identified by BLAST search using Arabidopsis RIN4 as query; the alignment was generated using CLUSTALW (https://www.genome.jp/tools-bin/clustalw). Two conserved motifs (PxFGxW and Y/FTxxF) in RIN4-like proteins are indicated by the red and blue open boxes, respectively.

**Figure S13.**
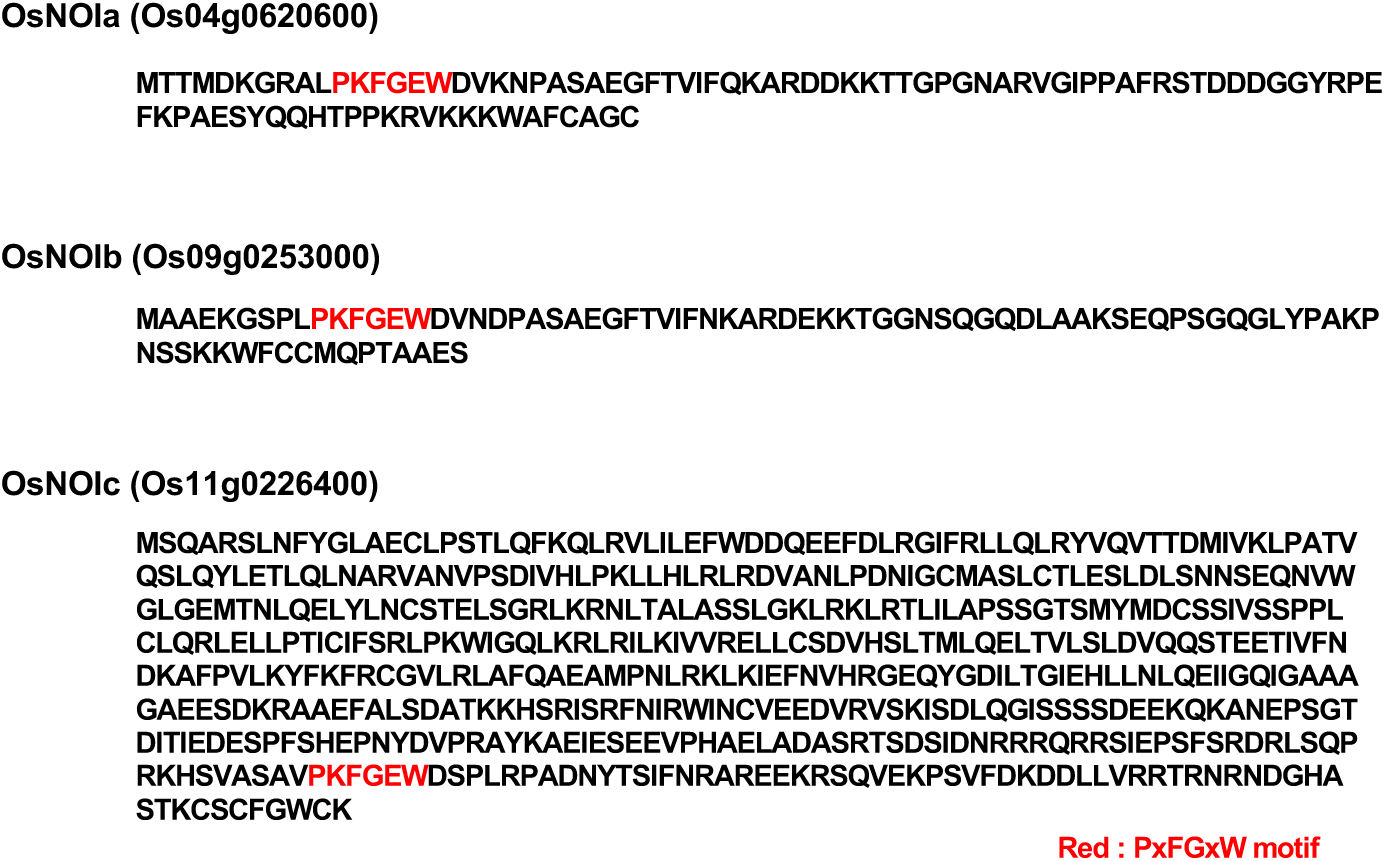
Amino acid sequences of rice PxFGxW motif-containing proteins that are not related to OsRIN4. Deduced amino acid sequences of rice PxFGxW motif–containing proteins used in this study. A BLAST search using Arabidopsis RIN4 as query identified three PxFGxW motif–containing proteins that are not related to OsRIN4s. These proteins were designated as OsNOIa (Os04g0620600), OsNOIb (Os09g0253000), and OsNOIc (Os11g0226400). The PxFGxW motif is marked in red text.

**Figure S14.**
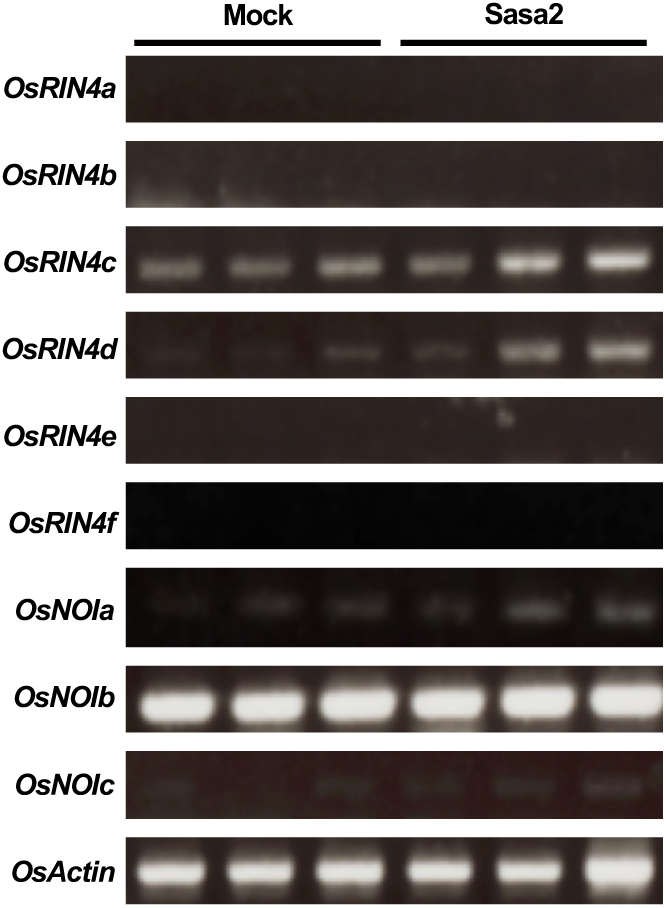
Expression of *RIN4-like* genes and other genes encoding NOI-containing proteins in rice leaves. RT-PCR analysis was used to confirm the expression of *RIN4-like* genes and other genes encoding NOI-containing proteins in rice leaves. Leaves of the rice cultivar Hitomebore were mock-inoculated or inoculated with rice blast fungus (Sasa2); leaves were harvested at 3 days post-inoculation, and total RNAs were extracted and used for RT-PCR using specific primers. *OsACTIN* served as a positive control.

**Figure S15.**
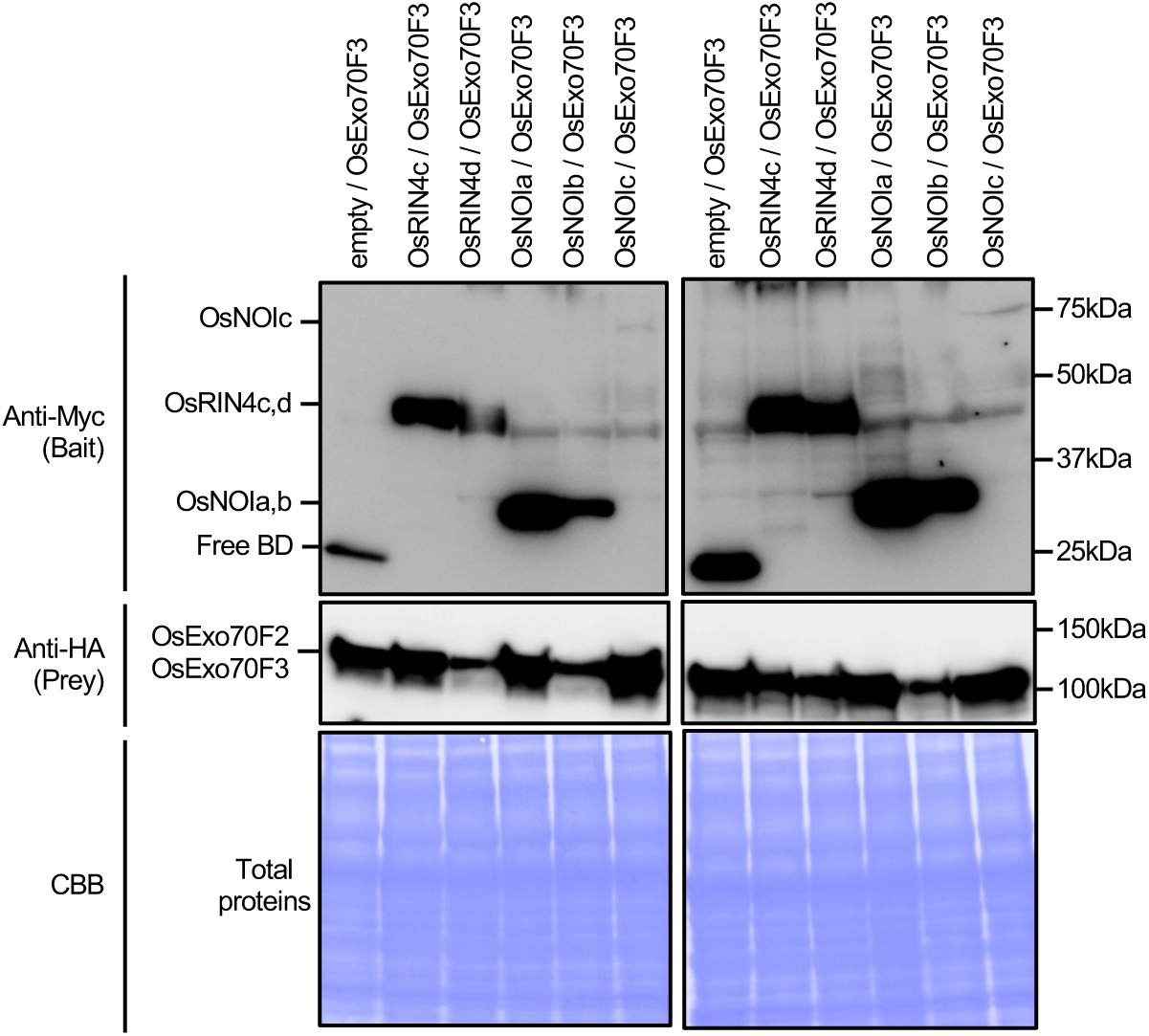
Confirmation of accumulation of OsExo70F2, OsExo70F3, and PxFGxW motif– containing proteins in yeast cells used in the Y2H assay. Immunoblot analysis showing the accumulation of proteins with the expected molecular weight in yeast cells used in Y2H assay testing the interaction between OsExo70F2 or OsExo70F3 and rice PxFGxW motif–containing proteins (Fig. 1F). The presence of proteins encoded by the bait and prey vectors was probed using anti-Myc and anti-HA antibodies, respectively, and is shown in the upper (bait) and lower panels (prey). The expected position of proteins is indicated at left. A CBB-stained proteins are shown below as loading control.

**Figure S16.**
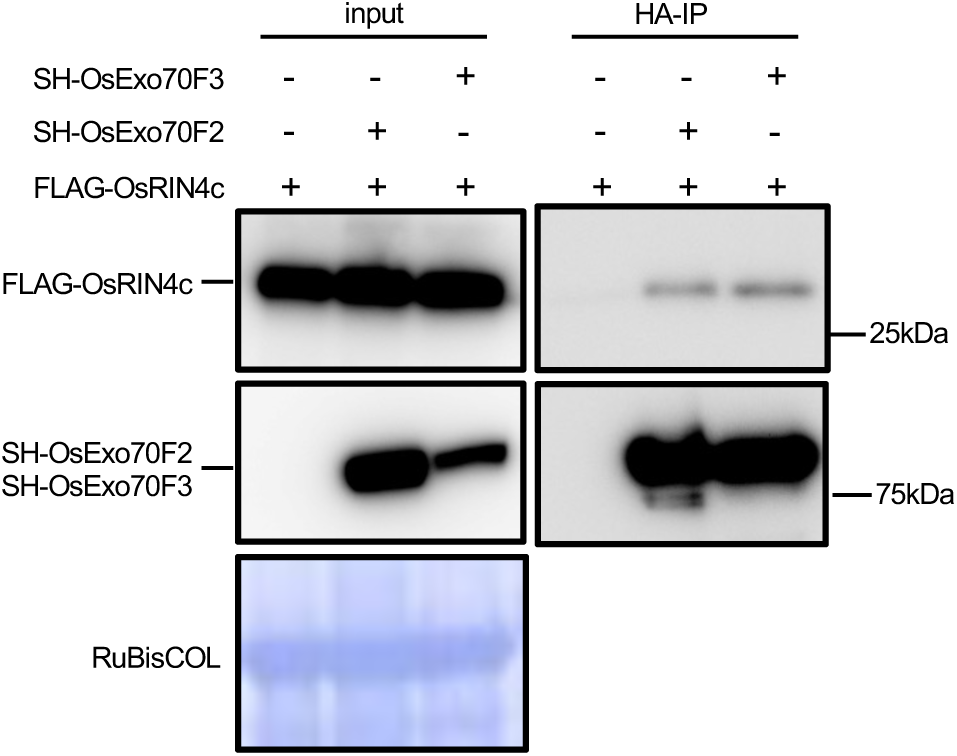
Confirmation of the interaction between OsRIN4c and OsExo70F2 or OsExo70F3 by co-immunoprecipitation assay. Constructs encoding FLAG-tagged OsRIN4c and/or StrepII-HA (SH)-tagged OsExo70F2/F3 were infiltrated into *N. benthamiana* leaves in the indicated combinations. The infiltrated leaves were sprayed with 30 μM dexamethasone, and total proteins were extracted at 2 days after the treatment for co-immunoprecipitation with anti-HA antibodies. CBB-stained RuBisCOL (Rubisco large subunit) are shown below as loading control of input proteins.

**Figure S17.**
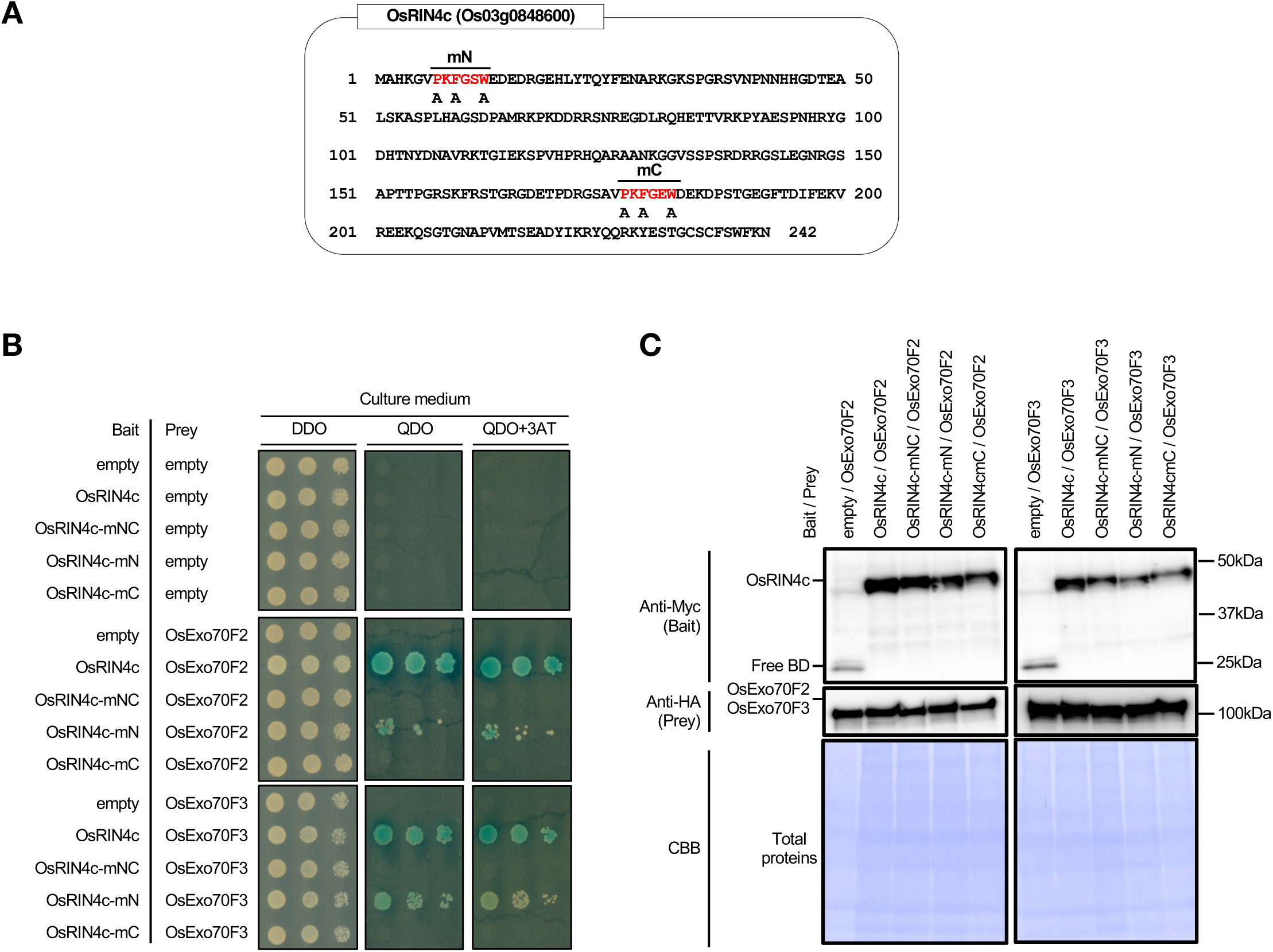
The C-terminal NOI core motif of OsRIN4c is involved in its interaction with OsExo70F2 and OsExo70F3. (A) Positions of the mutations in the mN and mC variants of the PxFGxW motif (red letters) of OsRIN4c. (B) Y2H assay to test the interaction of OsExo70F2 or OsExo70F3 and intact OsRIN4c or OsRIN4c carrying the mN and/or mC mutations. Ten-fold serial dilution of positive yeast transformants were spotted onto DDO medium as control and on QDO medium and QDO medium + 10 mM 3AT to test interactions. Empty vector (empty) was used as negative control. Bait and prey combinations are listed at left. (C) Immunoblot analysis showing the accumulation of proteins with the expected molecular weight in yeast cells used in the Y2H assay testing the interaction between OsExo70F2 or OsExo70F3 and OsRIN4c. The presence of proteins encoded by the bait and prey vectors was probed using anti-Myc and anti-HA antibodies, respectively, and shown in the upper (bait) and lower panels (prey). The expected position of proteins is indicated at left. A CBB-stained proteins are shown below as loading control.

**Figure S18.**
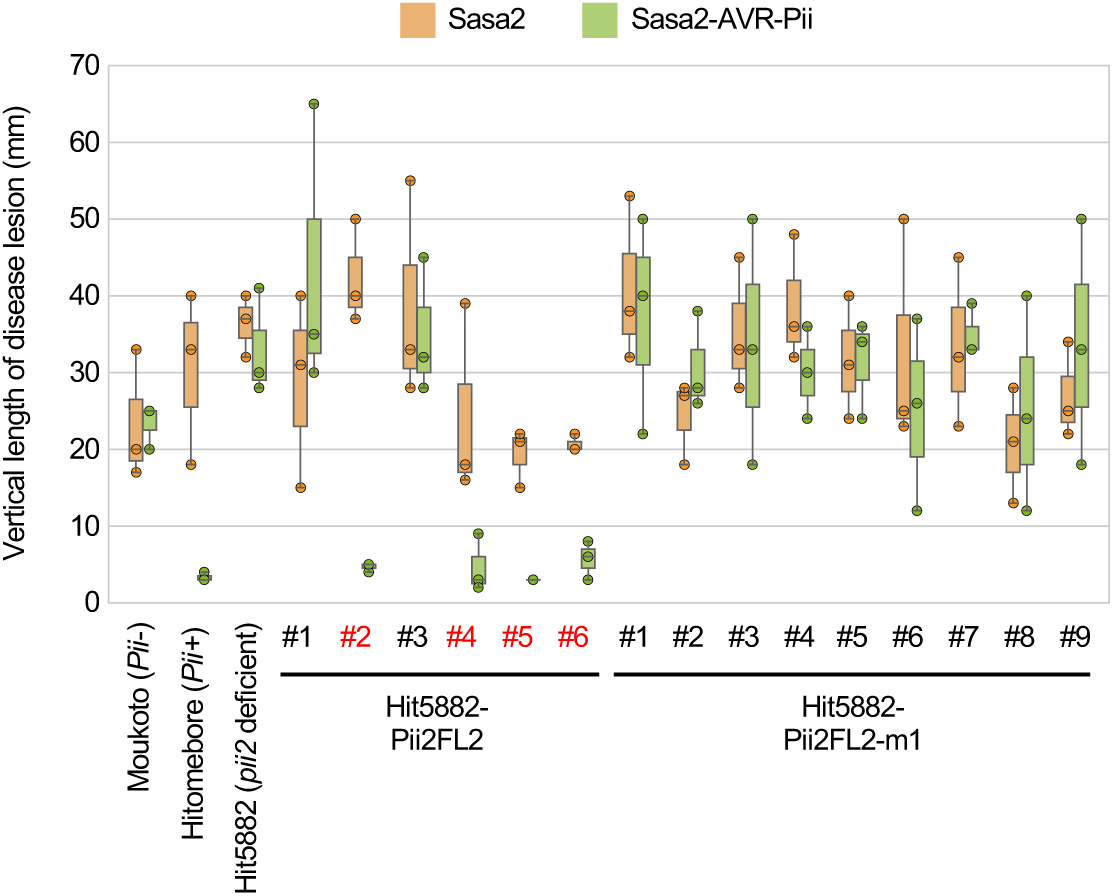
The Pii2 variant carrying the m1 mutation does not induce resistance against AVR-Pii. Length of disease lesions in the rice lines tested. Boxplots were illustrated from the values of three inoculated spots of rice blast fungus Sasa2 (lacking *AVR-Pii*) and Sasa2–AVR-Pii (with *AVR-Pii*) per line. The ttransgenic lines showing *Pii*-mediated resistance are marked by red letters.

**Figure S19.**
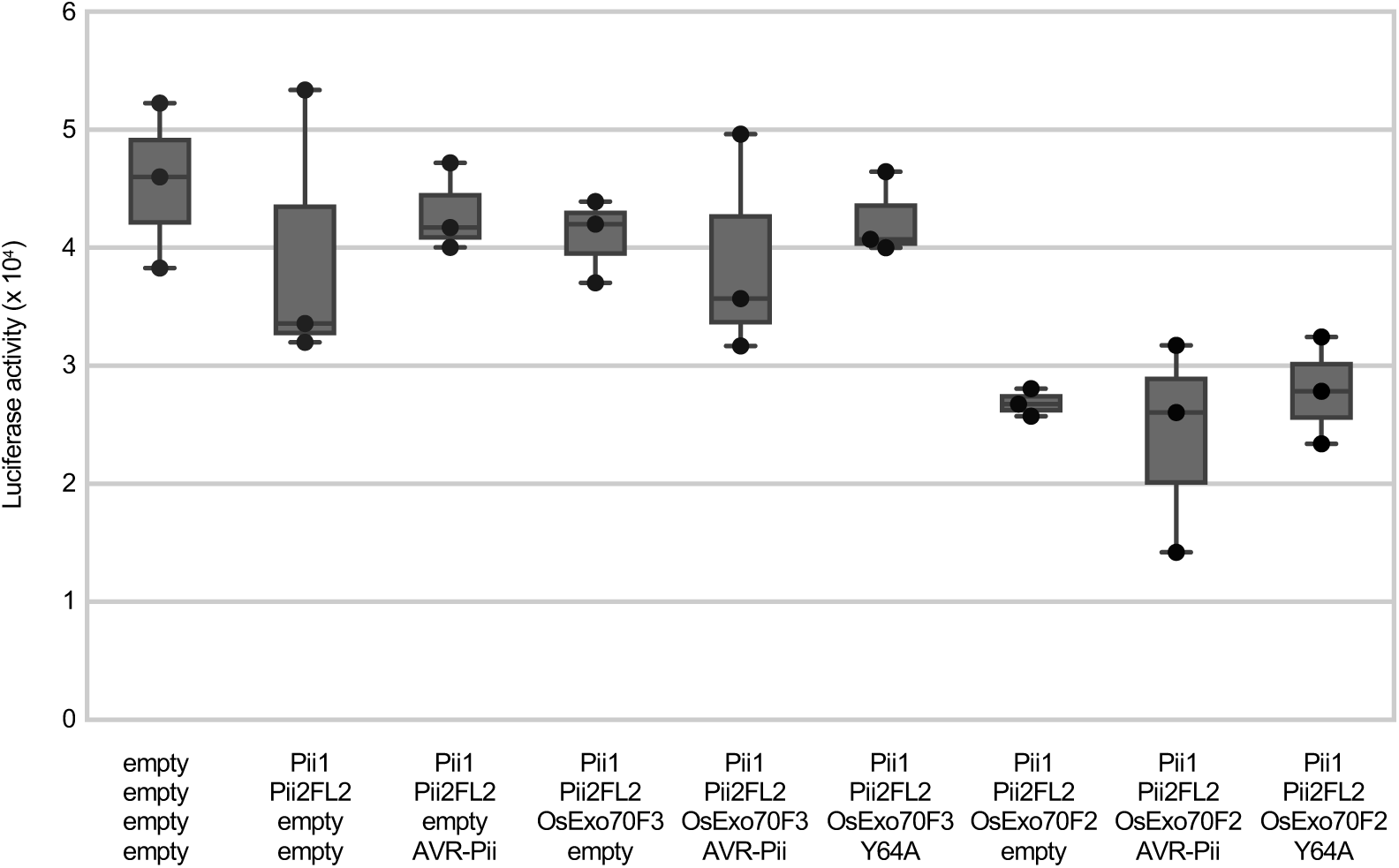
Viability of rice protoplasts transfected with various Pii components. Cell viability was estimated as described previously (Yoshida et al., 2009). *Firefly luciferase* gene was expressed under the control of Cauliflower mosaic virus 35S promoter in ice protoplasts (Oc cell). Several expression vectors of the components of *Pii*-mediated resistance (Pii1, Pii2FL2, OsExo70F2, OsExo70F3, FLAG-AVR-Pii and FLAG-AVR-Pii^Y64A^) were also co-expressed. The combinations of expression vectors are indicated bellow. FLAG-AVR-Pii and FLAG-AVR-Pii^Y64A^ are indicated as AVR-Pii and Y64A, respectively. Protoplasts were collected at 20 hours post-transfection, and luciferase activity was measured. Boxplots were illustrated from the values of three technical replicates.

**Figure S20.**
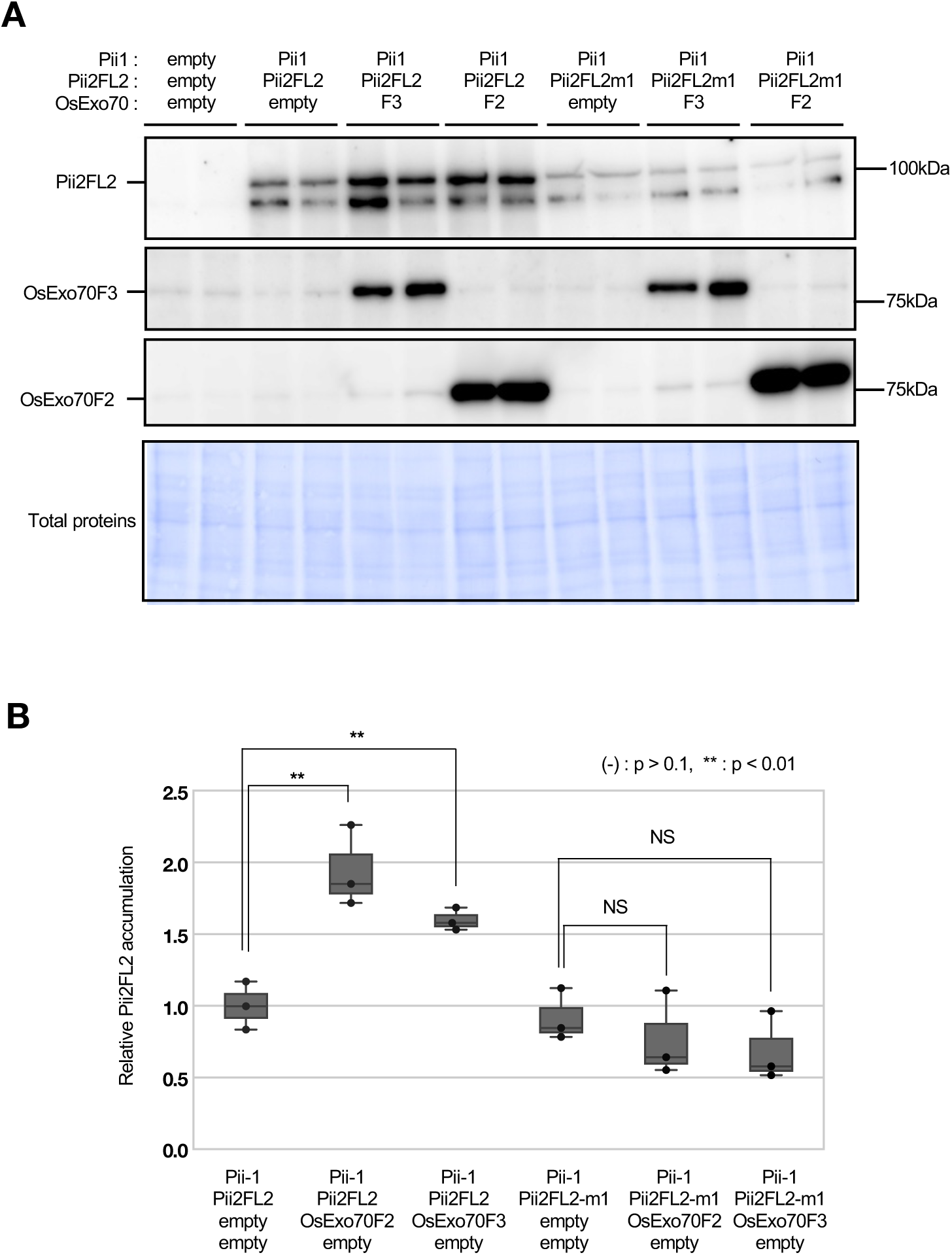
Effects of the m1 mutation on OsExo70F2- or OsExo70F3-mediated stabilization of Pii2FL2 in rice Oc protoplasts. (A) Representative data for the abundance of Pii2FL2 and Pii2FL2-m1 when their encoding constructs were co-expressed with or without *OsExo70F2* or *OsExo70F3* in rice Oc protoplasts. (B) Quantification of the abundance of Pii2FL2 and Pii2FL2-m1 shown in (A). Protein band intensity was quantified using ImageJ (https://imagej.net/ij/). Double asterisks represent significant differences (*t*-test; *P*<0.01), and NS indicates no significant differences in Pii2FL2 accumulation levels. Boxplots were illustrated from the values of three technical replicates.

**Figure S21.**
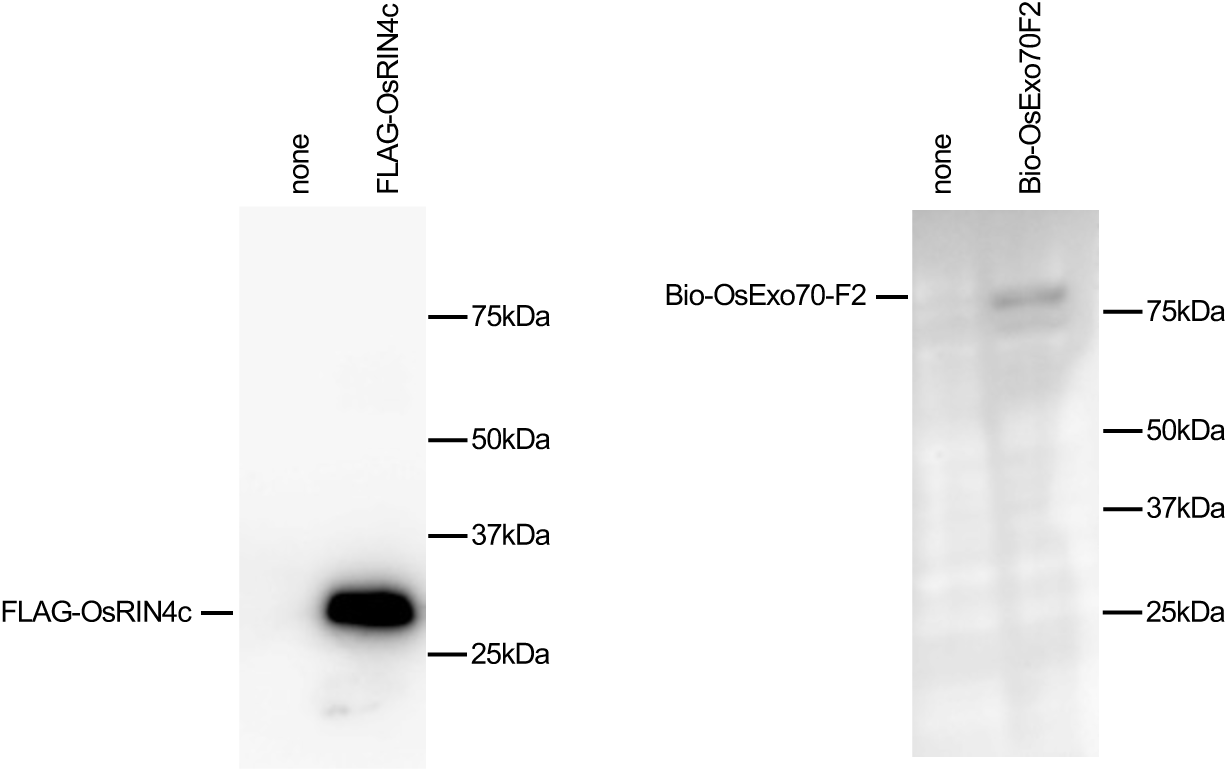
Confirmation of proteins synthesized by WGE used for AlphaScreen. (A) Biotinylated and FLAG-tagged proteins for AlphaScreen were synthesized in the WGE system. Biotinylated OsExo70F2 and FLAG-tagged OsRIN4c (FLAG-OsRIN4c) were used for the AlphaScreen (Fig. 5B). Biotinylated (right) and FLAG-tagged (left) proteins were detected using streptavidin and anti-FLAG antibodies, respectively.

